# Down-regulation of *Drosophila* Glutactin, a cholinesterase-like adhesion molecule of the basement membrane, impairs development, compromises adult function and shortens lifespan

**DOI:** 10.1101/2022.11.30.518525

**Authors:** Pedro Alvarez-Ortiz, Shawna Guillemette, Rachel Humphrey, Bryan A. Ballif, Jim O. Vigoreaux

## Abstract

Basement membranes (BM) play fundamental roles in morphogenesis and tissue maintenance in multicellular organisms. Glutactin is a BM protein that belongs to the Cholinesterase-Like Adhesion Molecules (CLAMs) protein family. In *Drosophila* embryos, Glutactin has been shown to outline internal organs and to play a role in synapse formation. Here, we report that Glutactin is broadly expressed in BM surrounding most vital tissues of the larva and the adult, and within the larval muscle sarcomere. Ubiquitous RNAi driven down-regulation of Glutactin expression (*Tub*>*Glt-RNAi*) resulted in pronounced impairments in larval and adult locomotor behavior, reduced oviposition, and shortened lifespan. Muscle-specific down-regulation of Glutactin resulted in reduced larval crawling speed indicating a secondary function for Glutactin independent of BM expression. *Tub*>*Glt-RNAi* pupa showed abdominal scars, suggestive of defects in histoblast nest expansion and replacement of larval epidermal cells, and a high mortality rate at eclosion. Surviving adults showed a range of morphological and physiological defects including excess melanization and pigmentation, incomplete rotation and duplication of the genitalia, and abnormal heart morphology and contraction. Insofar excess melanization is symptomatic of internal tissue damage, we propose that Glutactin is essential for the mechanical stabilization of the BM and for its ability to withstand internal stresses.

## 1. Introduction

The basement membrane (BM) is a specialized scaffold of extracellular matrix (ECM) found on the basal side of all polarized epithelia and surrounding muscle, adipose and neural tissues^1,2^. BM fulfills a variety of functions during morphogenesis, tissue remodeling, and tissue repair^3^. Conditions that compromise the structural integrity of the BM can lead to cell growth deregulation and tumor formation^4^. BM proteins are restricted to metazoans, suggesting these proteins played an important role in the evolution of the Kingdom Animalia.

The BM helps protect skeletal muscle from the strain and stresses of contraction^5^. The dystroglycan-dystrophin complex connects BM proteins such as Laminin, Agrin, and Perlecan to the intracellular cytoskeleton (for review, see^6^). Other common BM proteins include Collagen (type IV, XV and XVII) and Perlecan^7^. Additionally, BM also contain members of a family of adhesive proteins known as CLAMs (cholinesterase-like adhesion molecules) that are widespread in metazoans and are involved in the development of different tissues^8^. Invertebrate CLAMs include Glutactin Gliotactin, Neurotactin, and Neuroligin. Only Neuroligin is expressed in vertebrates. Among CLAMs, only Glutactin lacks a transmembrane domain^8^. With the exception of Glutactin, other members of this family have been shown to share a conserved function in cell adhesion and junction formation and are known to be expressed in a wide range of tissues throughout different stages of development^8^. Characterization of the expression and function of Glutactin will shed light on the evolution of the CLAMs family of proteins.

In *Drosophila*, a single gene encodes Glutactin, an ~117 kDa BM protein (1026 amino acids) that migrates as a 155 kDa protein on SDS-PAGE possibly due to its high negative charge resulting from a large number of acidic residues (52 glutamates) and four O-linked-sulfated tyrosine in its C-terminus^9^. The N-terminal, ~ 500 amino acid region has 34% identity with the acetylcholinesterase (AChE) domain from humans, but the AChE-like domain in Glutactin has no enzymatic activity due to the absence of a catalytically essential serine residue^9^. The C-terminal, ~400 amino acid region of Glutactin has no homology to known protein domains. Separating the N-terminal and C-terminal regions is a stretch of 13 consecutive threonine residues. The cholinesterase-like N-terminal region demonstrates adhesive properties in a cell aggregation assay^10^. No function has been assigned to the threonine stretch or the C-terminal region.

Glutactin was originally isolated from *Drosophila* Kc cells and later found in embryonic hemocytes, the primary secretory source of Glutactin and other BM proteins^11^. Immunofluorescence staining of whole mount *Drosophila* embryos detected Glutactin in the BM outlining internal organs including the gut, brain, and the nerve cord^9^. Subsequent studies found that Glutactin acts as a repulsive molecule that inhibits synapse formation to muscle M12 during embryonic development (stage >16)^12^. Despite these studies, very little else is known about Glutactin. More than two decades after its discovery, Glutactin remains a chronically understudied protein of the BM^2^.

Here, we seek to gain additional insight into the expression and functional roles of Glutactin in developing and adult *Drosophila*. Specifically, we aim to establish if this unique member of the CLAMs family functions post-embryogenesis and whether its presence is required for normal morphogenesis and function of adult organs. We report that Glutactin is widely expressed in *Drosophila* larval and adult tissues and that RNAi-mediated knock down of Glutactin results in morphological and motility defects, a high mortality rate at eclosion, and a shortened lifespan. We also report that Glutactin is expressed in muscle where it also contributes to normal motility.

## 2. Results

### 2.1. Glutactin is essential for normal lifespan

To determine if Glutactin plays a role during adulthood, we used the ubiquitously expressed Tubulin-Gal4 driver (*Tub*>) to down-regulate the expression of Glutactin by RNAi and compared the lifespan of *Tub*>*Glt-RNAi* flies to *Glt-RNAi*/+, and *Tub*>/+ flies. The RNAi construct used in these studies has a calculated target specificity (s_19_) of 0.997^13^. Knockdown in Glutactin expression was assessed by western blots (Supplementary Figure S3-3). Due to the temperature sensitivity of the ectopic expression of Gal4 in *Drosophila*^14–16^, we conducted the experiments at two different temperatures, 29°C and 25°C, to compare phenotypic effects at high and medium penetrance, respectively. Figure 1 shows a high mortality of *Tub*>*Glt-RNAi* flies in comparison to the controls *Tub*>/+ and *Glt-RNAi*/+, where 50% of *Tub*>*Glt-RNAi* adult flies died by day 4 vs day 38 for the parental lines. In addition, the maximum lifespan of *Tub*>*Glt-RNAi* flies was shorter, 37 days at 29°C compared to 50 days for control strains. A similar pattern was observed at 25°C where the median lifespan for *Tub*>*Glt-RNAi* flies was ≤4 days and >50 days for the parental lines (data not shown). In contrast, we found that constitutive gain-of-function of Glutactin (*Tub*>*UAS-Glt*) resulted in a lethal phenotype at the pre-instar stages, suggesting that Glutactin levels above normal are even more detrimental. This early lethality was partially rescued by reducing Glutactin expression with RNAi. Flies carrying two transgenes *Tub*>*Glt-RNAi;UAS-Glt* not only completed development but also overcame the early adult mortality evident in *Tub*>*Glt-RNAi* (Supplementary Figure S1-1). These studies demonstrate an important requirement of Glutactin for development and adulthood.

**Figure 1.**
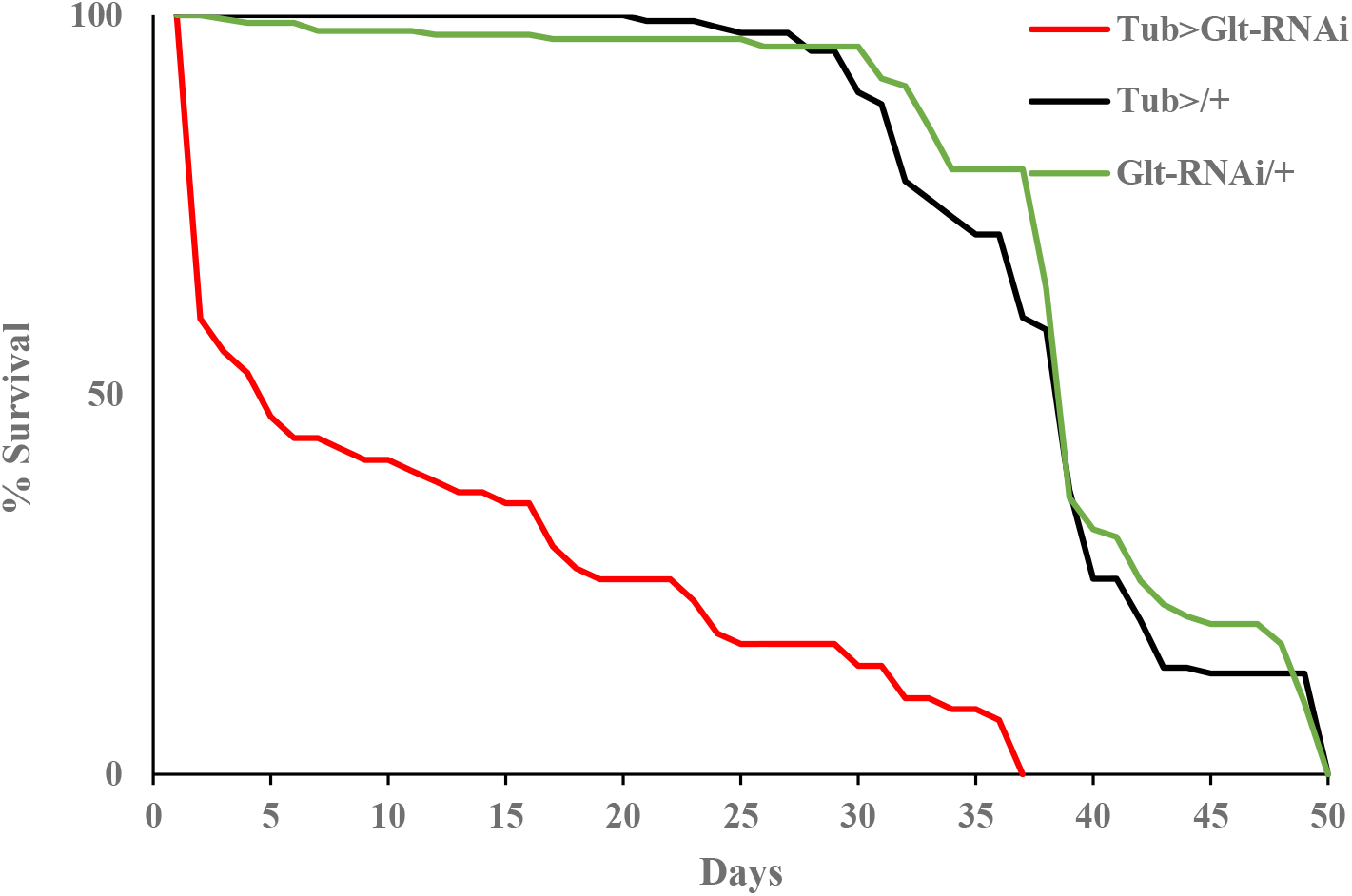
RNAi down regulation of Glutactin shortens lifespan. Down-regulation of Glutactin resulted in high mortality during the first four days after eclosion. Maximum lifespan for *Tub*>*Glt-RNAi* flies (red line, n = 70) was 37 days, compared to 50 days for both *Glt-RNAi* /+ (green line, n = 128) and for *Tub*>/+ (black line, n = 192). Experiment was performed at 29°C.

The majority of *Tub*>*Glt-RNAi* adults lost locomotory functions abruptly, within one to three days after eclosion, and exhibited trembling and jerky movements (Supplementary video 1). These flies walked sporadically and did not fly or jump. A smaller subset of *Tub*>*Glt-RNAi* adults died within the first day after eclosion before exhibiting walking, jumping, or flying ability. Further evidence that the expression of Glutactin is essential for adult viability comes from studies documenting the genotypic ratio of the progeny from the cross *Tub-Gal4/TM6B* x *Glt-RNAi*. The expected ratio is 50% *Glt-RNAi*/+; *TM6B*/+ and 50% *Glt-RNAi*/+; *Tub-Gal4*/+ while the observed ratio was 77% *Glt-RNAi*/+; *TM6B*/+ and 23% *Glt-RNAi*/+; *Tub-Gal4*/+ at 29°C (n=2,945), and 74% *Glt-RNAi*/+; *TM6B*/+ and 26% *Glt-RNAi*/+; *Tub-Gal4*/+ at 25°C (n=1,143). The similar but slightly more penetrant phenotypes observed at 29°C vs 25°C is consistent with previous studies showing a greater expression of Gal4 at 29°C^14,15^.

Members of the CLAMs family, in particular neuroligins, are expressed strongly in neurons. To determine if neuronal expression of Glutactin is responsible for adult mortality and loss of gross locomotory control, we used the pan-neural driver elav-Gal4 to increase and decrease the expression of Glutactin. Neither up-regulation nor down-regulation of Glutactin affect lifespan and the adult flies have none of the visible phenotypes observed with the tubulin down-regulation (Supplementary Figure S1-2).

### 2.2. Down-regulation of Glutactin causes morphological abnormalities in adult structures

*Tub*>*Glt-RNAi* adult flies exhibited a wide range of morphological abnormalities. While each individual phenotype was rare (Supplementary Table S2.1), collectively these abnormalities were common features among surviving *Tub*>*Glt-RNAi* flies. The abdomen of adult flies was most commonly affected, ranging from a completely collapsed abdomen (Figure 2B-C) to differences in morphology, pigmentation, and segmentation (Figure 2 and Supplementary Figure S2). In some cases, female genitalia was characterized by an extended and exposed ovipositor (Figure 2H) with a darkly pigmented or necrotic tip (Figure 2I). Male genitalia was also affected by down-regulation of Glutactin, exhibiting incomplete rotation (Figure 2K) or duplication (Figure 2L) of the genital disk. Internal organs such as the heart tube were also affected as evidenced by misaligned pericardial cells that produced abnormal peristaltic movements (Supplementary video 2). The thorax occasionally showed darker pigmentation of varying shades (Supplementary Figure S2, K-L). Flies exhibiting a darker pigmentation throughout most of their bodies often showed no movements and died within 22-24 hours.

**Figure 2.**
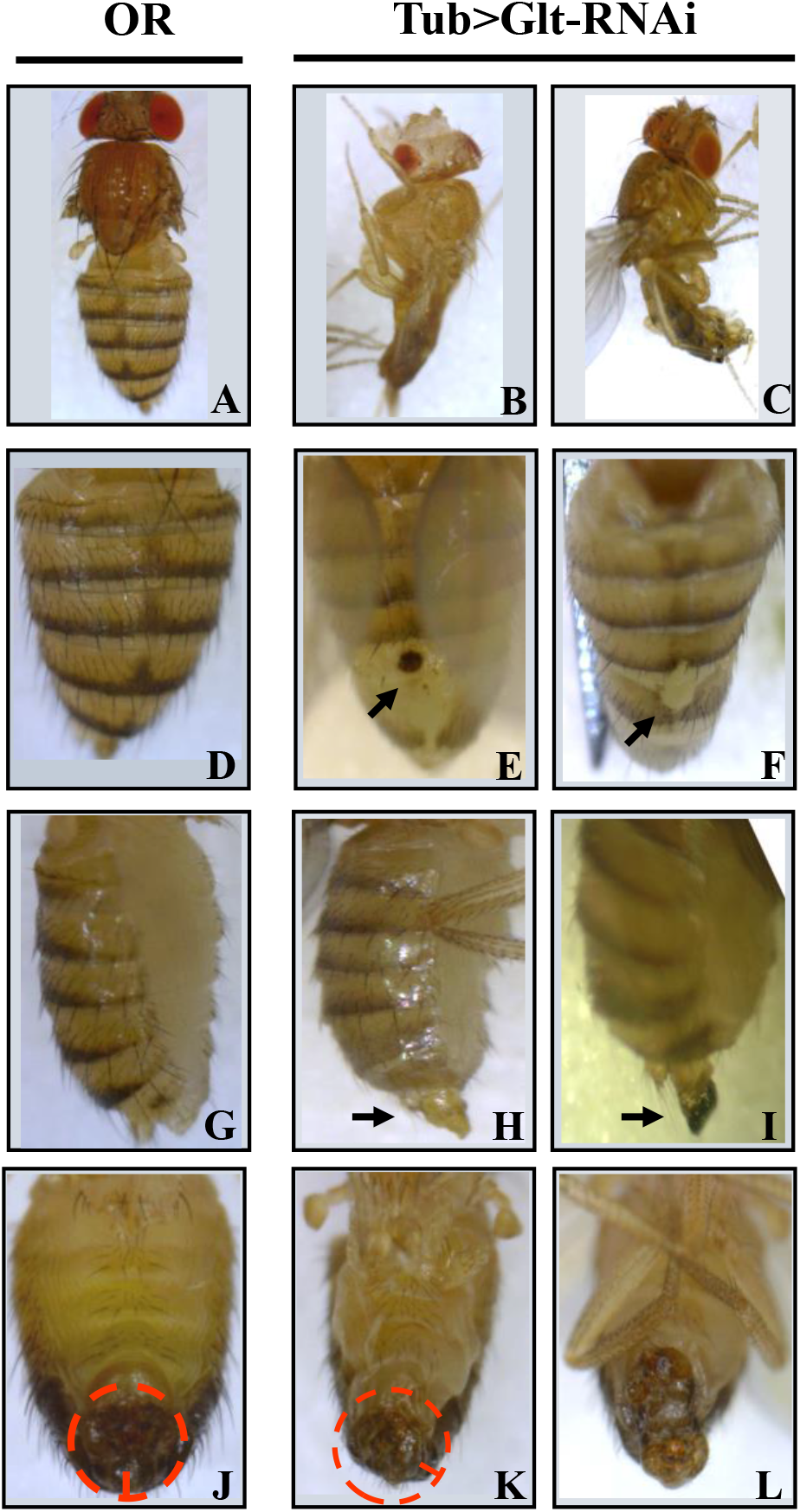
Morphological changes in the abdomen associated with Glutactin down-regulation by RNAi. Oregon R flies (OR) are shown for direct comparison of the morphological changes (panels A, D, G, J); all other panels are *Tub*>*Glt-RNAi* flies. In most panels, wings were clipped to allow better viewing. B and C, examples of recently eclosed flies showing a common collapsed abdomen phenotype. Panel E and F, examples of abnormal pigmentation on the dorsal side of the abdomen (arrows). Panel H and I, side view of female abdomen with extended ovipositor (arrows), which occasionally showed dark pigmentation or necrosis at the tip (I). Panel K and L, ventral view of male abdomen with genitalia showing incomplete rotation (K) and duplication (L).

The diversity of morphological phenotypes observed in *Tub*>*Glt-RNAi* adult flies suggest that expression of Glutactin is associated with many tissues and organs, most notably the reproductive system, circulatory system (heart), and possibly the digestive system. None of above phenotypes have been observed in flies from the combined down- and up-regulation line *Tub*>*Glt-RNAi;UAS-Glt*, further evidence that the absence of Glutactin is largely responsible for the observed defects.

### 2.3. Glutactin is expressed broadly in adults

The expression of Glutactin in adult flies was assessed in frozen sections of two to three days old OR flies stained with mouse monoclonal antibody (mAb) C7906-50. The validation of mAb C7906-50 specificity against Glutactin is presented in Supplement 3 and consisted of western blots of proteins from dissected tissues and body parts (Figure S3-1), pull-down assays followed by protein identification by mass spectrometry (Figure S3-2), and western blots of proteins from larva of different genetic strains (Figure S3-3). Glutactin expression was detected throughout the adult body, most prominently surrounding the abdominal intestinal tract, the reproductive organs, tubular muscles of leg and thorax, and the brain (Figure 3 and 4, Supplementary Figure S3.1 and Supplement 4).

**Figure 3.**
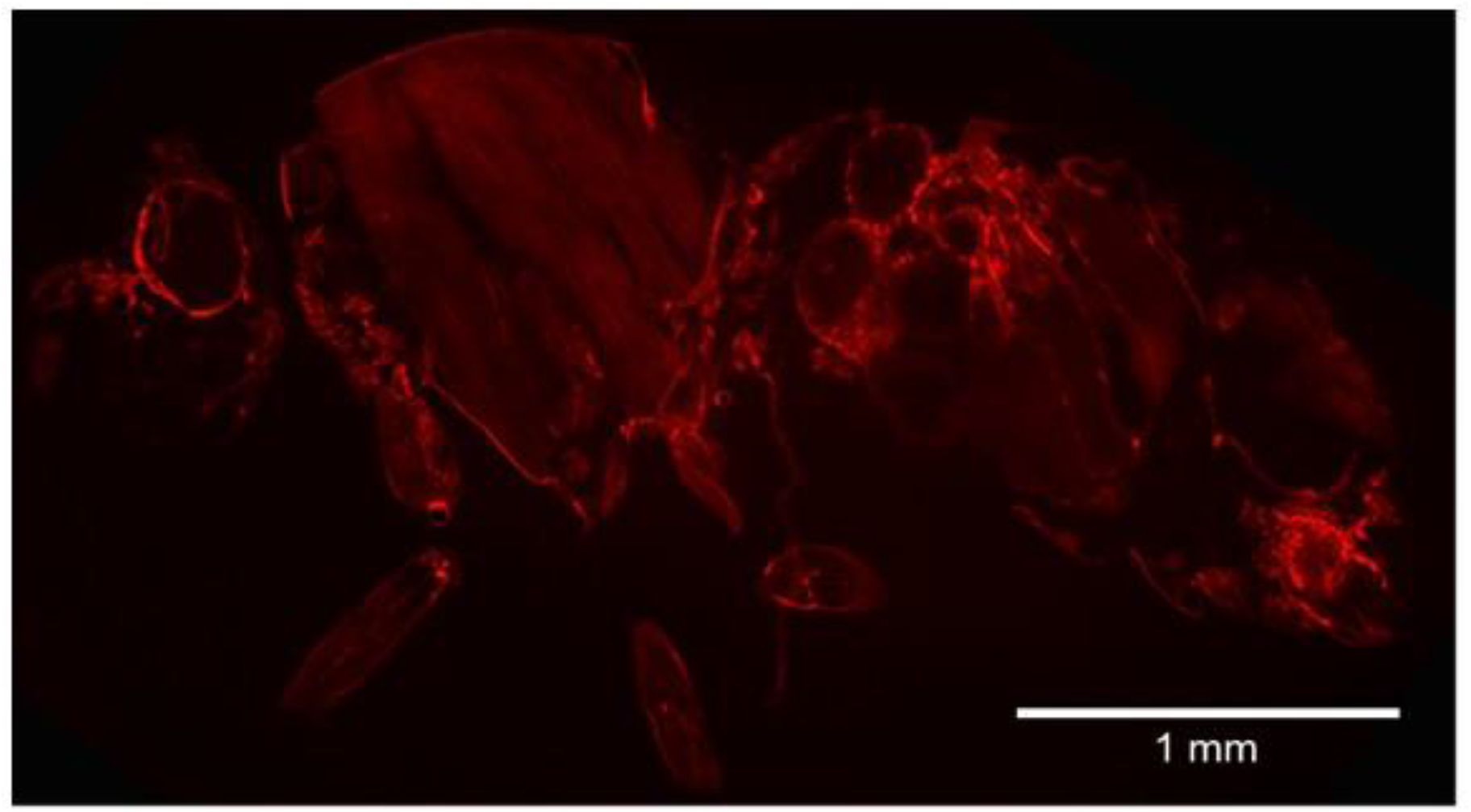
Glutactin expression is widespread in adult flies. Shown is a sagittal section through an adult female stained with anti-Glutactin. Anterior is to the left, dorsal at the top. Glutactin expression is prominent in the abdominal intestinal tract, in the genitalia, and in the tubular muscles of the leg and the thorax (see Supplement 4 for additional images).

**Figure 4.**
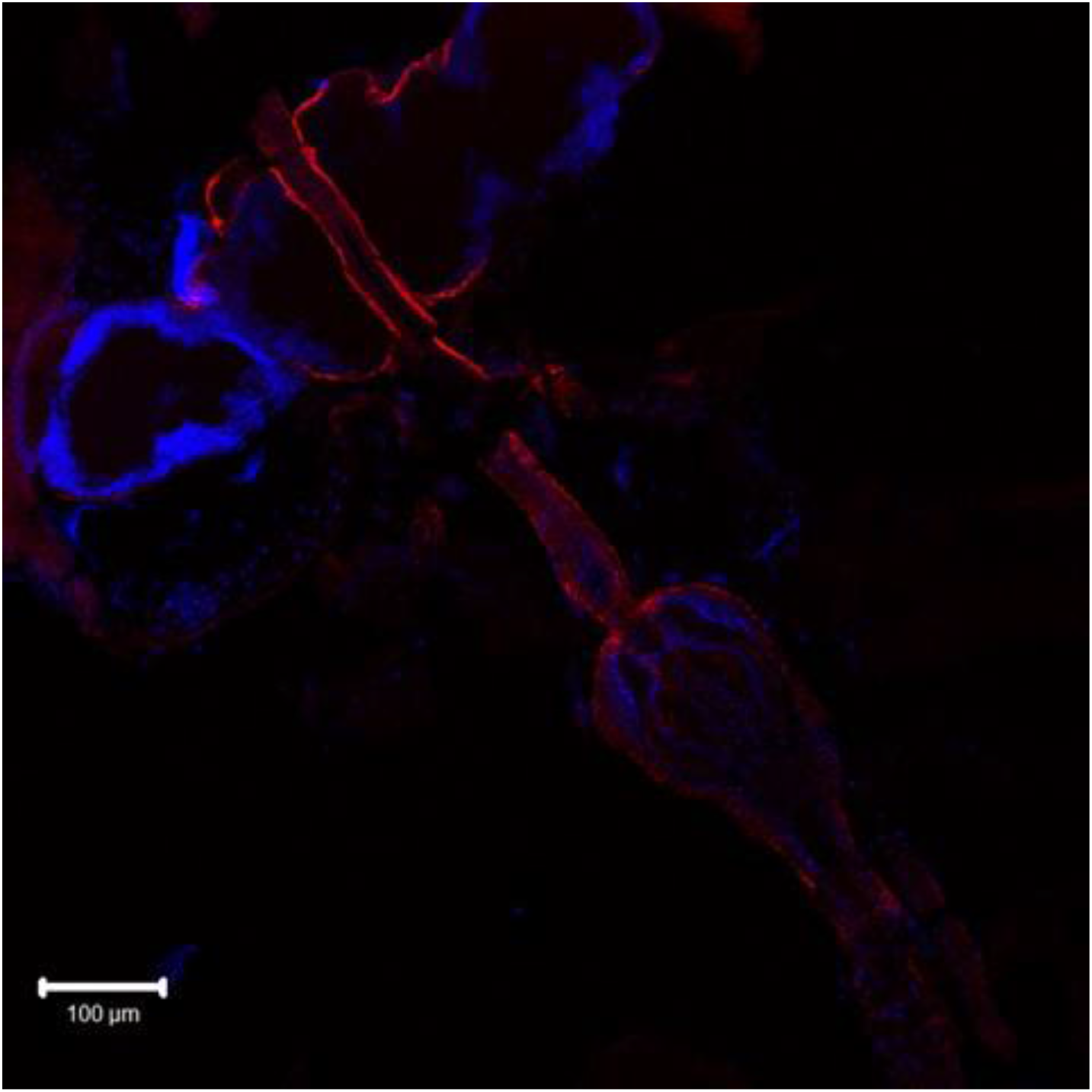
Glutactin expression in the anterior region. Shown is a horizontal section of an adult female double immunostained with anti-Glutactin (red) and DAPI (blue). Glutactin is detected surrounding the brain and foregut (esophagus and proventriculus). Anterior is top left. Faint autofluorescence is seen in the eye region.

With respect to the gut and intestinal tract, Glutactin expression appeared limited to the peripheral visceral muscles (Supplementary Figures S4.1: A-C; S4.2: A-B; S4.3: D), the proventriculus (Figure 4 and Supplementary Figure S4.2B), and the rectum (Supplementary Figure S4.3: C). With respect to the reproductive organs, Glutactin was strongly expressed in both females (ovaries, oviduct, uterus, and vagina; Supplementary Figure S4.3: A-C) and males (testis, accessory gland, ejaculatory bulb, and penis; Supplementary Figure S4.4: A-F). In the thorax we detected Glutactin expression surrounding the direct flight muscles and, less prominently, surrounding the indirect flight muscles (Supplementary Figure S4.5, A-D). Glutactin was also detected surrounding the tubular leg muscles and, additionally, along transverse stripes resembling the stereotypical striation pattern of skeletal muscles (Supplementary Figure S4.6). In summary, Glutactin appears to be associated with most, if not all, vital tissues and organs of the adult fly.

### 2.4. Glutactin down-regulation affects egg laying

Given the strong expression of Glutactin associated with the female reproductive system (Supplementary Figure S4.3) we conducted experiments to determine if down-regulation of Glutactin affects egg laying. Many of the *Tub*>*Glt-RNAi* flies died during the first three to four days and for these experiments we used virgin females that survived more than 6 days. When comparing the number of eggs laid during nine days by *Tub*>*Glt-RNAi* virgin flies to virgin females of the parental lines *Tub*>/+ and *Glt-RNAi*/+, we found that down-regulation of Glutactin resulted in a significant reduction in oviposition (Figure 5). There was also a significant difference between the two parental lines: *Tub*>/+ females laid half as many eggs (10 ± 2/day) than *Glt-RNAi*/+ females (21 ± 3/day) (Figure 5). This would indicate that the severe reduction in eggs laid by *Tub*>*Glt-RNAi* (2 ± 0.4/day) cannot be completely ascribed to the absence of Glutactin. However, we also noticed that the proportion of females that fail to lay eggs was 3x-4x higher in *Tub*>*Glt-RNAi* compared to the two parental lines.

**Figure 5.**
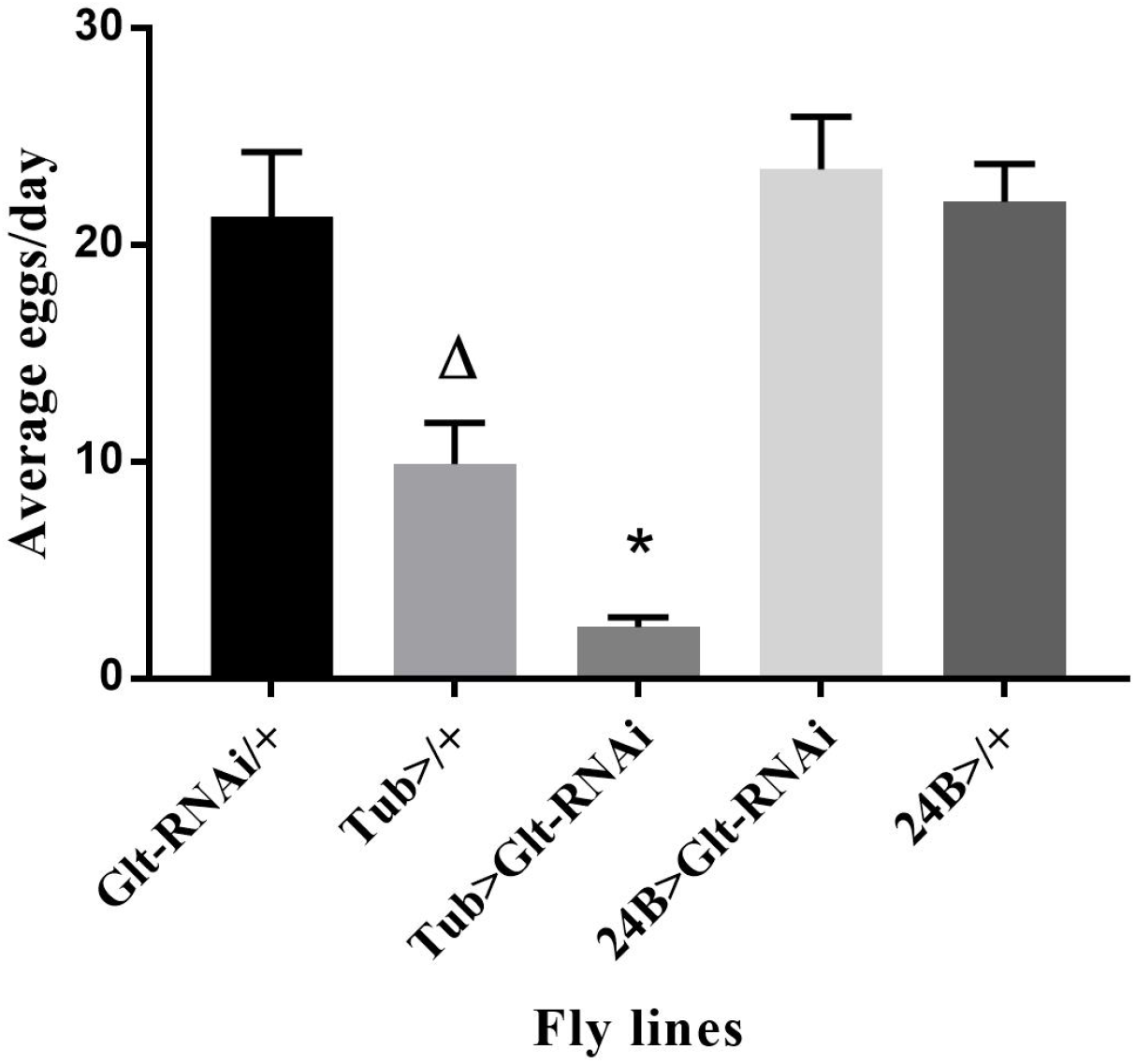
Oviposition is compromised with Glutactin down-regulation by RNAi. Virgin females were individually kept in separate vials and allowed to lay eggs for a period of 15 days, with counting of eggs beginning at day seven. Down-regulation of Glutactin decreases the average number of eggs laid per day. Bars indicate standard error. Asterisk (*) denotes significant difference from controls, *Glt-RNAi*, and *Tub*>; (p<0.001), and from samples *24B*>*Glt-RNAi* and 24B>; (p<0.001) by One-way ANOVA. Δ denotes significant difference from *Glt-RNAi*(p<0.001). N = 9 for all samples.

Previously, we showed that down-regulation of Glutactin with the *Tub*> driver resulted in an exposed ovipositor with dark pigmentation or necrosis at the tip (Figure 2). To determine if ovipositor exposure was due to problems in the ovipositor muscles, which may lead to reduced oviposition and sterility, we tested the effect of down-regulation of Glutactin using the mesoderm specific Gal4 driver *24B*>. There was no difference in the number of eggs laid by *24B*>*Glt-RNAi* females (23 ± 3/day) when compared to females from control lines *Glt-RNAi*/+ (21 ± 3/day) and *24B*>/+ (22 ± 2/day) (Figure 5).

### 2.5. Glutactin is required for progression through pupal development and for eclosion

The cross *Tub*>/TM6B x *Glt-RNAi* resulted in a larger than expected number of dead pupa at 29°C as well as at 25°C. To determine if down-regulation of Glutactin was responsible for the pupal lethality, we used *Tubby*, a marker on the TM6B balancer chromosome, to discriminate between the two progeny genotypes, *Tub*>*Glt-RNAi* and *Glt-RNAi*/+; TM6B/+. At 29°C nearly all of the animals that die during early (i.e., before eye pigmentation) and late (i.e., after eye pigmentation) pupal stages are *Tub*>*Glt-RNAi*, while at 25°C all of the dead pupa are *Tub*>*Glt-RNAi* (Table 1). To determine if down-regulation of Glutactin is solely responsible for the mortality, we conducted an additional control experiment by crossing *Tub*>/TM6B to *w^1118^*. Presence of the *Tub*> chromosome resulted in ~4% pupal mortality at 29°C, significantly below the levels seen among *Tub*>*Glt-RNAi* pupa. Mortality at eclosion is also highly skewed, with >99% of the dead pharate-adults being *Tub*>*Glt-RNAi* (Table 1).

**Table 1.** Down-regulation of Glutactin interferes with pupal development and eclosion.

| Genotypes | Temp<br>°C | Pupa |  | Pharate-Adult |
| --- | --- | --- | --- | --- |
|  |  | Early<br>%<br>(counts) | Late<br>%<br>(counts) | %<br>(counts) |
| Glt-RNAi/+; TM6B/+ | 29 | 4.7<br>(15) | 10.6<br>(42) | 0.4<br>(1) |
| Tub>Glt-RNAi |  | 95.3<br>(305) | 89.4*<br>(355) | 99.6<br>(277) |
| Glt-RNAi/+; TM6B/+ | 25 | 0 | 0 | 0 |
| Tub>Glt-RNAi |  | 100<br>(21) | 100<br>(201) | 100<br>(224) |

Numbers indicate percentage and numbers of flies (in parenthesis) whose development is arrested at early and late pupal stages, and pharates who fail to eclose. *In a separate experiment we determined that ~4% *Tub-Gal4*/+ pupae are arrested at late stages when grown at 29°C.

Many of these flies were unable to emerge completely from the pupal case because part of their abdomen remained glued to the pupal case (Figure 6). Manual removal of these flies out of the pupal case revealed a severe abdominal lesion surrounded by necrotic-like tissue at the point of contact with a semi-transparent sheath that covered the whole body (Figure 6C-F).

**Figure 6.**
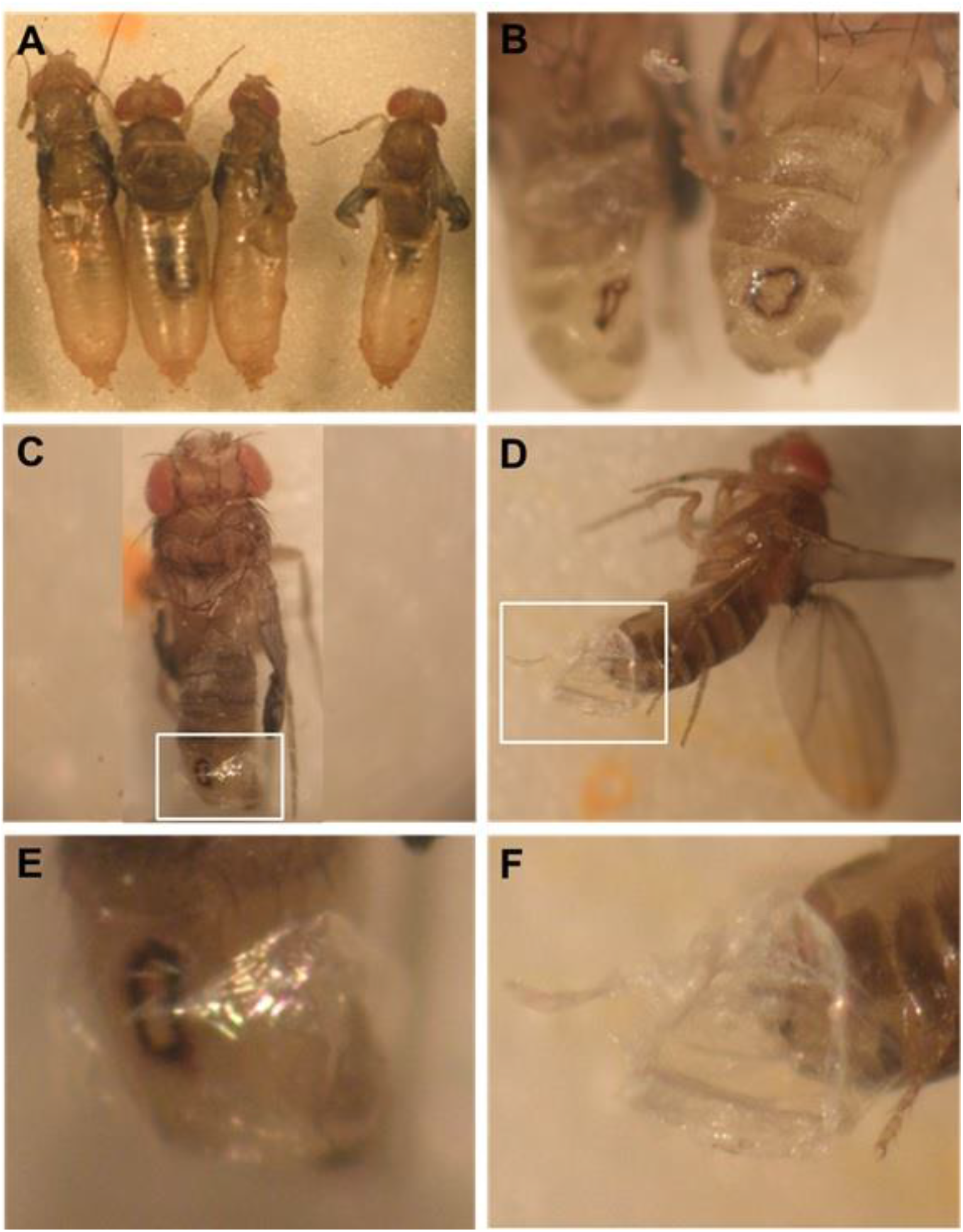
RNAi down-regulation of Glutactin expression affects eclosion. Panels A-F, examples of *Tub*>*Glt-RNAi* flies that failed to eclose from the pupal case. Panel B, higher magnification of the abdomen of two of the flies in A after removal from the pupal case. Note lesion surrounded by necrotic tissue; this area remained attached to the pupal case, likely preventing eclosion. A transparent sheet that wrapped the abdomen remained attached at the lesion site (Panel C-D). Panel E-F, higher magnification of boxed region in panel C-D respectively.

Down regulation of Glutactin produced many morphological defects in developing pupa, most prominently in the abdomen. A common phenotype was open scars on the dorsal side, with different degrees of penetrance (Figure 7 and Supplementary Figure S5.1 C-E). Another phenotype detected in pupa and dead pharate-adults was the presence of an off-white material present within the cuticle in the anterior end of the pupa (Supplementary Figure S5.1A-E). While less common, we also observed a similar dried white material covering the head and thorax of living adults (not shown).

**Figure 7.**
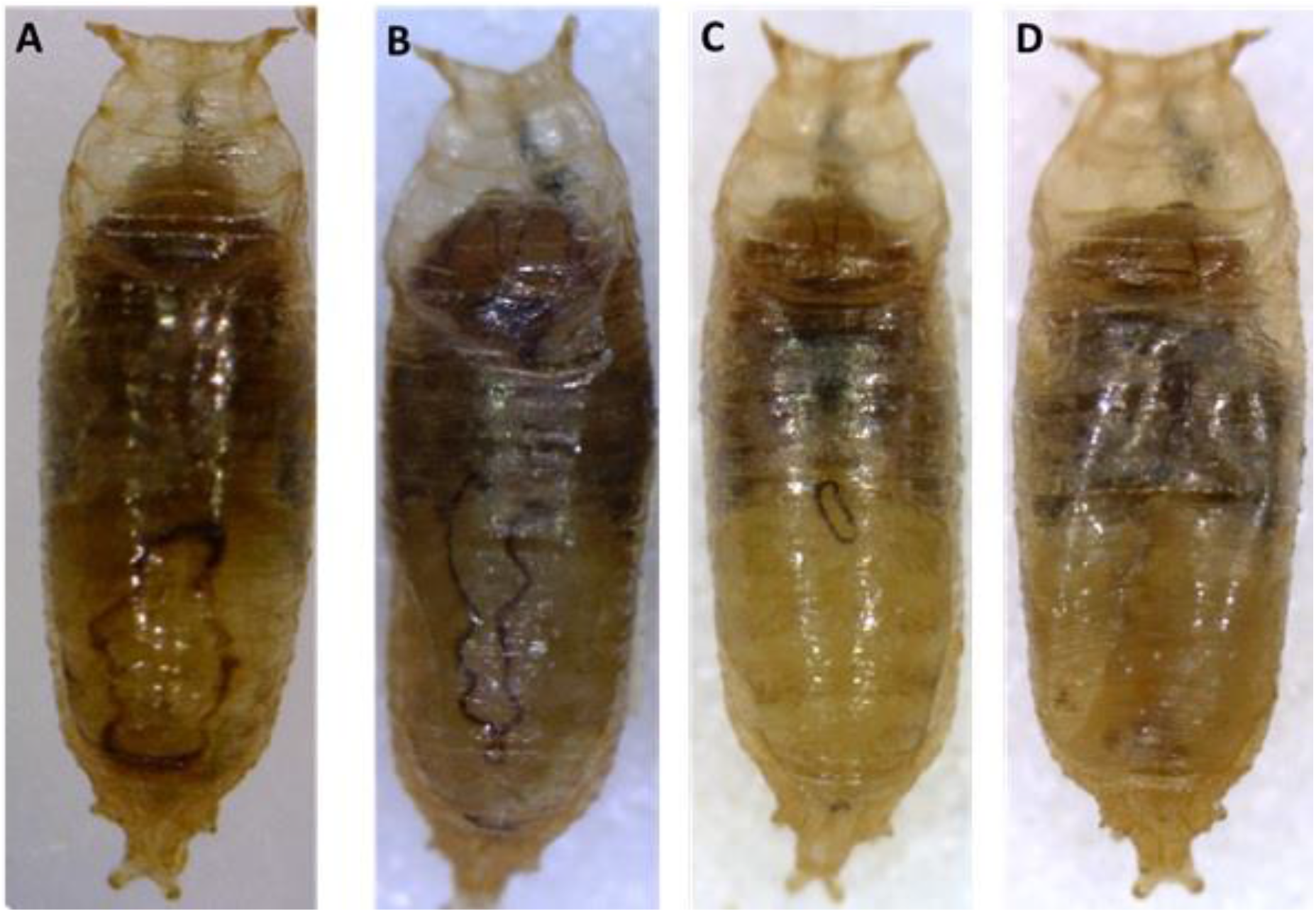
Down-regulation of Glutactin affects pupal development. Panels A-D, dorsal view of *Tub*>*Glt-RNAi* pupa. In all panels anterior is up. Panels A to C, examples of different severity of scars in the abdominal region of developing pupa. Panel D, abnormal pigmentation phenotype in the abdominal region of developing pupa.

### 2.6. Glutactin down-regulation affects larval motility

*Tub*>*Glt-RNAi* pupa are commonly found closer to the food than *Glt-RNAi*/TM6B pupa, an indication that *Tub*>*Glt-RNAi* larva do not crawl too far before metamorphosis. To determine if this is due to a motility defect, we measured the crawling speed of individual larva^17^. *Tub*>*Glt-RNAi* larva crawling speed (0.044 ± 0.001 cm/s) (N = 87) was significantly slower than both of the parental strains, *Tub*>/+ (0.067 ± 0.002 cm/s; n=42) (p<0.001) and *Glt-RNAi*/+ (0.081 ± 0.002 cm/s; n=38) (p<0.001). The crawling speed of *Glt-RNAi*/+ larva is comparable to that reported earlier for Canton-S and Oregon-R strains, 0.081 cm/s^18^. The slower crawling speed detected in the *Tub*>/+ control, while not statistically significant from that of *Glt-RNAi*/+ control, could be due adverse effects from the expression of Gal4^16^.

To gain insight into how down-regulation of Glutactin affects larval motility, we examined its expression by immunostaining of whole mount larva.

### 2.7. Glutactin expression in larval basement membrane and muscle

Glutactin is expressed broadly throughout the body wall of third instar larva (Fig. 8A). To determine if Glutactin localizes to the BM, we used a *trol-GFP* fly strain as a reference. *Trol* encodes DPerlecan, the *Drosophila* homolog of the BM protein Perlecan^19^. DPerlecan expression during embryonic development has been reported in the BM covering the channels of the CNS, the BM surrounding the gut, dorsal median cells, and dorsal muscle attachments sites^20^, a distribution that is comparable to what has been described for Glutactin^9,21^. In the larva, DPerlecan expression has been detected in the BM surrounding imaginal discs^22,23^. As seen in Fig. 8 A-H, DPerlecan, detected with an anti-GFP antibody, co-localizes with Glutactin throughout most of the larval body wall, specifically in the BM surrounding the muscles. Given this co-localization, we then tested if the expression of Glutactin and DPerlecan is inter-dependent.

**Figure 8.**
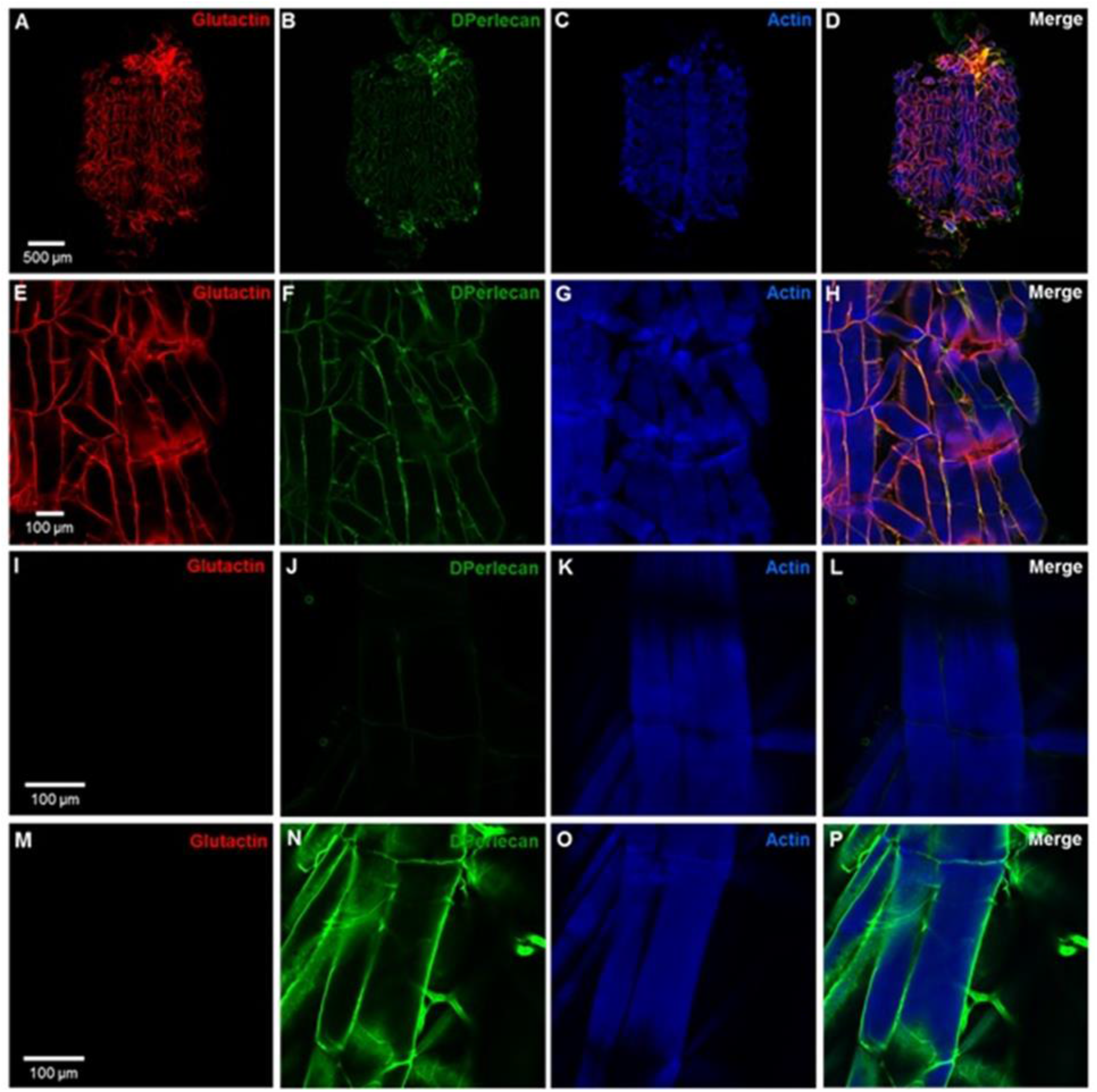
Co-localization of Glutactin and DPerlecan expression along the body wall muscles of 3^rd^ instar larva. In all the images anterior is at the top. A-D. Confocal images of larva fillets (*trol-GFP*) labeled with anti-Glutactin (red), anti-GFP (DPerlecan; green), and phalloidin (actin; blue). E-H. Higher magnification confocal images of the right side abdominal segments 3 and 4. The merged image (H) shows co-localization of Glutactin and DPerlecan (yellow-orange). I-L, larva (*trol-GFP*; *Glt-RNAi*/+) fillets showing muscle 7 and 6 labeled with secondary antibodies and phalloidin only. No signal was detected in I and the faint signal in J is from the endogenous GFP. M-P, down-regulation of Glutactin in (*trol-*GFP; *Tub*>*Glt-RNAi*). No Glutactin signal was detected (M) while DPerlecan signal remained relatively unchanged (N, P).

Down regulation of Glutactin did not affect expression of DPerlecan in the BM, as determined in a *trol-GFP; Tub*>*Glt-RNAi* fly strain (Fig. 8 M-P). However, the GFP staining appeared broader and more diffused than when Glutactin was present (*trol-GFP*, Fig. 8 F) suggesting that DPerlecan may require Glutactin for precise deposition along the BM or for proper expression. In contrast, BM deposition of Glutactin appeared to depend on the presence of DPerlecan. Female flies heterozygous for the DPerlecan hypomorphic *trol*^G0271^ allele, which contains a P-element insertion located in the 5’ UTR region ^24^, showed Glutactin and DPerlecan BM distribution and co-localization similar to that seen in OR flies (Fig. 9 A-D). The co-localization was also observed in hemizygous *trol*^G0271^ males, except in gaps in the BM region surrounding muscles 7 and 6 where both proteins were absent (Fig. 9 E-H). The absence of these BM proteins appeared to affect the immediately adjacent muscle, as evidenced by perturbations in the sarcomere striation pattern (Fig. 9 G-H). This myofibril phenotype was observed in different preparations but not in all areas devoid of DPerlecan.

**Figure 9.**
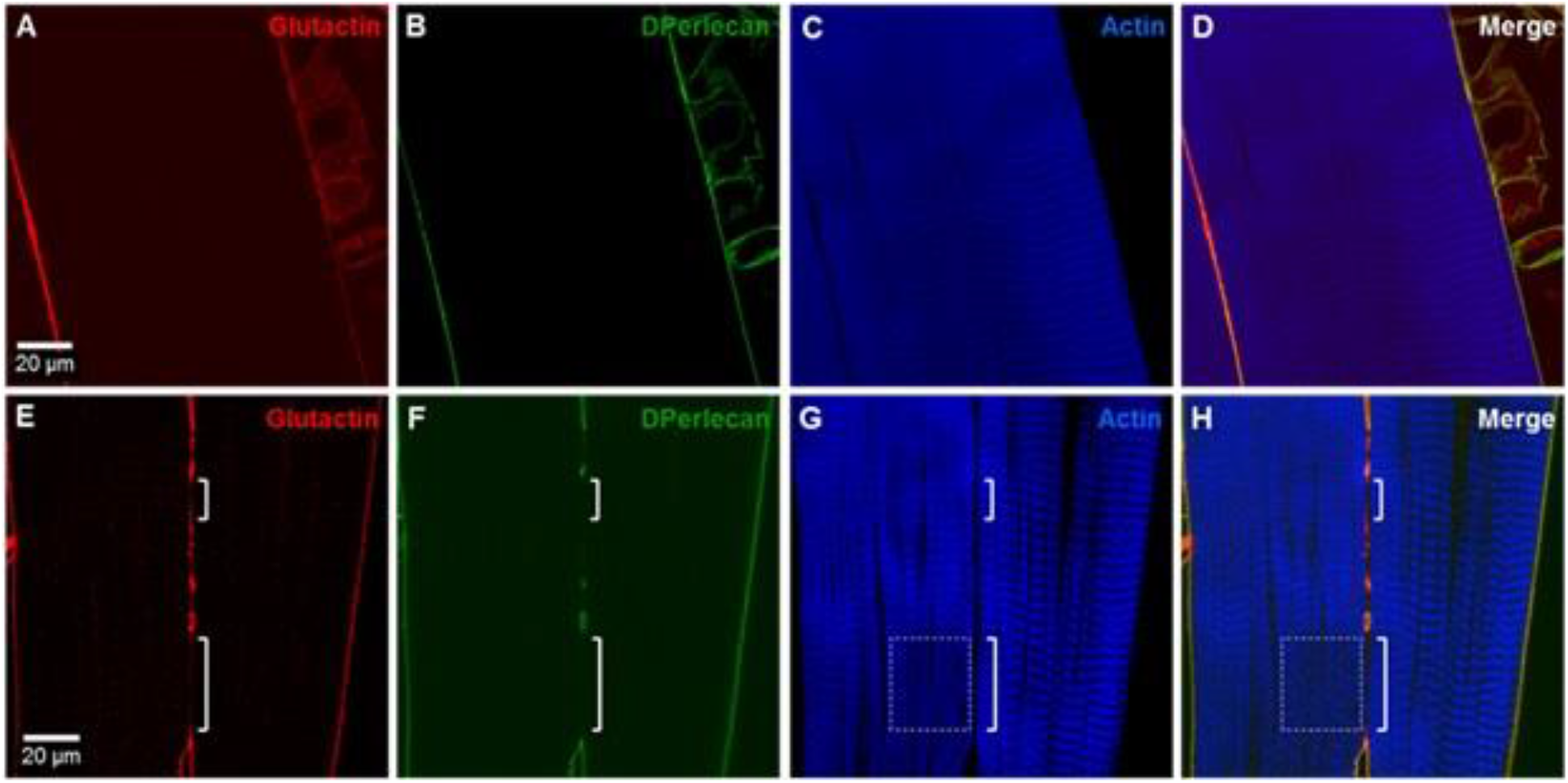
Glutactin is dependent on DPerlecan for its localization to the basement membrane. Images A-D are from OR larva while images E-H are from hemizygous larva of the hypomorph *trol*^G0271^. Panels A-D show muscle 6 and a small section of muscle 7 (lower left corner). In panel A and B Glutactin and DPerlecan are seen lining the muscle fiber. Note strong staining in the region between muscles 6 and 7 (lower left corner). Sarcomere striations appear in good registry across the fiber (C and D). Panel E and F show that when DPerlecan expression was reduced (green), localization of Glutactin (red) was affected, as evidenced by gaps in staining (brackets). Note that normal pattern of sarcomere striation was interrupted (box in G and H) in the region adjacent to large staining gap (lower bracket).

In addition to staining the larval BM, the anti-Glutactin antibody stained the body wall muscle weakly, in a pattern reminiscent of sarcomere striations detected in adult leg muscle (Supplementary Fig. S4.6). Glutactin co-localized with phalloidin in the I/Z region of the sarcomere (Figure 10, A-D). To confirm that Glutactin was indeed found within the muscle, we performed serial optical sectioning through 15μm of the nearly 20 μm thick tissue. The striation pattern and overlap with phalloidin staining was observed throughout the depth of the fiber (Figure 10, E-H).

**Figure 10.**
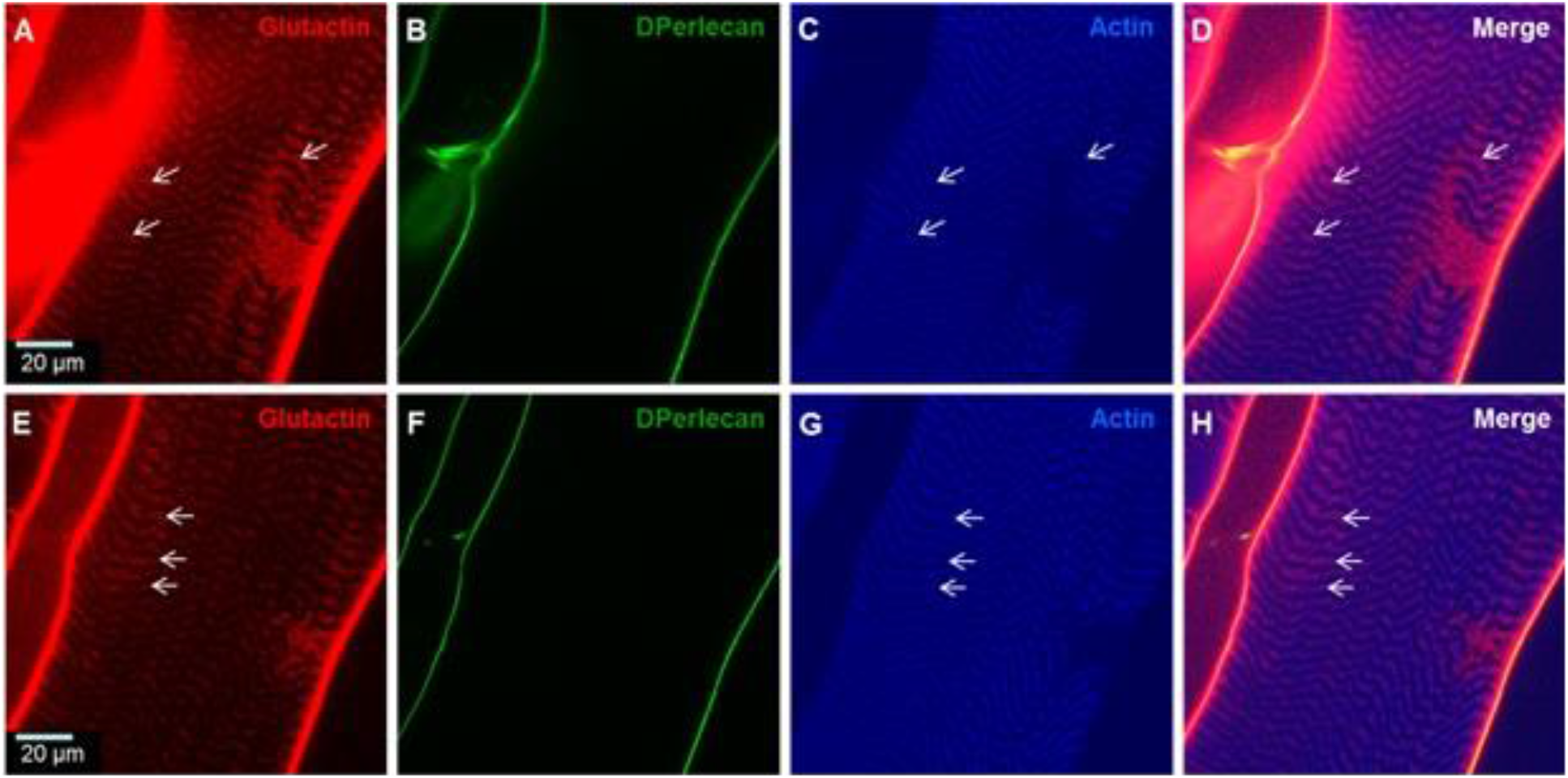
Glutactin overlays the I/Z region of the sarcomeres. In all the images anterior is at the top, muscle 7 is to the left and muscle 6 is to the right, A-D, muscles from larva (*Glt-RNAi*/+) stained for Glutactin (red), DPerlecan (green) and actin (blue). Glutactin staining pattern is rib-like and overlaps with phalloidin staining within the fiber and DPerlecan staining along the basement membrane (D). E through H are optical sections 15 μm below sections shown on A-D. Note that the Glutactin rib-like pattern overlapping the phalloidin staining (arrows) is still evident at this depth, as is the overlap with DPerlecan (yellow and orange).

Further evidence that Glutactin localizes to the I/Z region of the sarcomere comes from double immunostaining with anti-Glutactin and anti-Obscurin, an M line protein^25^. As seen in Fig. 11A-C, the Glutactin signal was detected intercalated between Obscurin-labeled M lines. Final confirmation that Glutactin was expressed in the larval muscle comes from RNAi down regulation of Glutactin using the mesoderm specific 24B-Gal4 driver. In *24B*>*Glt-RNAi* larva Glutactin was detected in the BM, however, the staining along muscle striations was significantly reduced (Fig. 11D). The reduced expression of Glutactin in the muscle did not appear to affect the expression and localization of Obscurin (Fig. 11E-F).

**Figure 11.**
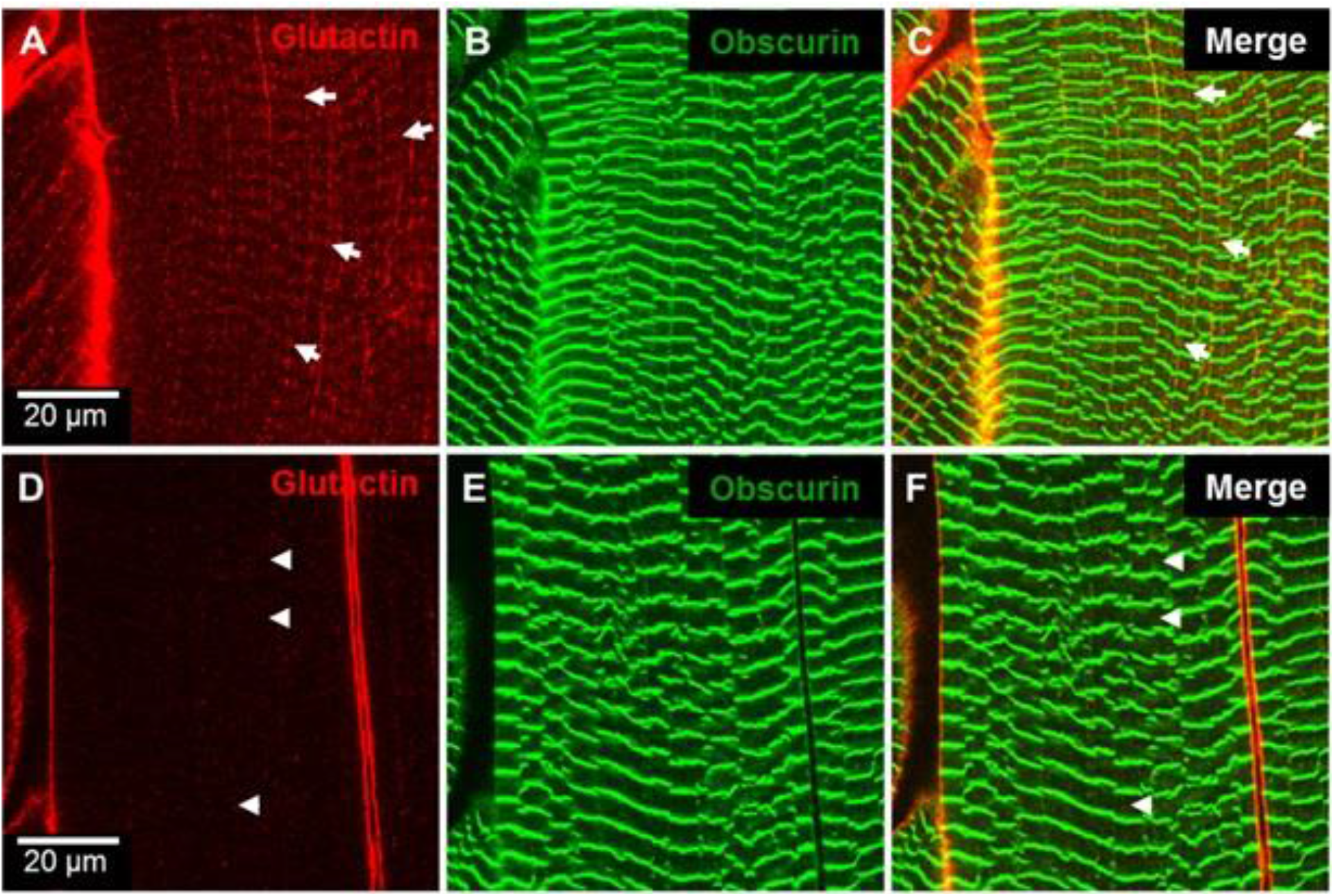
Down-regulation of Glutactin by muscle-specific driver prevents I/Z region localization. Optical sections showing muscle 7 of *24B*> larva (A-C) and *24B*>*Glt-RNAi* larva (D-F). Glutactin is in red and Obscurin in green. (A) Glutactin staining forms striations (arrows) along the length of the muscle. (C) shows Glutactin (arrows) localized in between striations of the M line marker, Obscurin. (D) Down regulation of Glutactin with the mesoderm specific driver *24B*> eliminates striations (arrowheads) in between M line marker Obscurin (F). BM staining remained intact (seen in the periphery on both sides of the muscle). (E and F) expression and sarcomere localization of Obscurin were not affected by down-regulation of Glutactin.

Given our earlier result that down regulation of Glutactin with the constitutive Tubulin driver caused a significant decrease in larval crawling speed, we next examined if mesoderm-specific down regulation with the 24B driver produced a similar effect. When compared to the parental strain *Glt-RNAi*/+, the *24B*> *Glt-RNAi* larva had a significantly slower crawling speed (0.072^±^0.003 cm/s (n=17) vs 0.085^±^0.003 cm/s (n=10); p<0.05). However, the crawling speed of *24B> Glt-RNAi* larva was not significantly different to that of the other parental line, *24B*>/+ (0.081^±^0.003 cm/s (n=10), p>0.05). Therefore, we tested another mesoderm specific driver, Mef2>, that has been reported to result in stronger expression than 24B>^26,27^. Our results revealed that *Mef2*> *Glt-RNAi* larva had a significantly slower crawling speed (0.068 ± 0.002 cm/s) (N = 35) compared to both parental strains, *Mef2*>/+ (0.081 ± 0.003 cm/s; n=24) (p<0.05) and *Glt-RNAi*/+ (0.100 ± 0.005 cm/s; n=27) (p<0.05).

## 3. Discussion and Conclusions

A major finding of this study is that Glutactin is essential for normal development and lifespan of *Drosophila*. Mis-regulation of Glutactin results in embryonic lethality (up-regulation) and in a variety of developmental, morphological, and physiological defects (down-regulation). Most notably, constitutive RNAi down-regulation of Glutactin resulted in defects in larval motility, high mortality during pupal development, at eclosion, and during early adulthood, and locomotory and reproductive defects in adults. These findings are consistent with numerous studies that have found important functional roles for BM in tissue organization and maintenance, morphogenesis, and normal organ function^2,28,29^. Our results, together with the finding that Glutactin is widely expressed in developing and adult flies, indicate Glutactin is a core component of the BM that enables its multiple functional roles. Additionally, we report that Glutactin expressed in larval muscle is required for normal locomotory function. Future studies should examine additional mutants of Glutactin to determine the extent to which distinct mutations contribute to the variety of phenotypes described in this study.

The shortened lifespan and high mortality during the first week of adult life when Glutactin is down-regulated indicate Glutactin is necessary for organ function in adults. Glutactin expression is most prominent in the abdomen where it is found surrounding the digestive track and the reproductive system in both males and females. Consistent with this finding is the observation that a large number of *Tub*>*Glt-RNAi* flies have abdomens with abnormal morphology suggesting that defects in the digestive system may be a primary cause of early mortality. An alternative, but not mutually exclusive, explanation is that developmental defects give rise to malfunctioning adult organs. The high incidence of pupal and pharate adult mortality, compared to a parental line (Table 1), is consistent with this notion, as is the frequent observation of deformations, duplications, and incomplete rotation of the genitalia. Mutants of the *Drosophila* ECM protein Tenectin exhibit reduced rotation of male genitalia, a phenotype consistent with a role of the ECM in left/right asymmetry^30^. Together, these results underscore the interplay between BM/ECM and epithelium-derived rotational forces in organ morphogenesis^31^. The anomalous genital structures observed in *Tub*>*Glt-RNAi* females can largely explain their greatly reduced rate of egg laying and their higher frequency of sterility. In addition, the BM is known to play an important, albeit not well understood, role in shaping the fly egg^32^. Thus, the decrease in the number of eggs laid by *Tub*>*Glt-RNAi* females could very well be a result of defects in egg elongation during oogenesis.

The remarkable variety of pupal phenotypes observed when Glutactin is down-regulated demonstrate the important role of Glutactin in developmental processes. Several of the abdominal defects resemble those described when the process of histoblast nest expansion and replacement of larval epidermal cells is interrupted by blocking the ecdysone signaling pathway^33^. Normally, circulating macrophagic hemocytes engulf and remove apoptotic larval epidermal cells to make room for the expanding histoblast nest. Failure to eliminate the larval epidermal cells prevents full expansion of the histoblast nest resulting in incomplete fusion of the abdominal epithelium. These pupa exhibit dorsal scars very similar to those seen in *Tub*>*Glt-RNAi* pupa suggesting that Glutactin may play a role in this process. The macrophagic hemocytes populate the BM of the abdominal epidermis and the absence of Glutactin may interfere with their positioning or ability to engulf the larval epidermal cells. Furthermore, hemocytes are known to express and secrete Glutactin, along with other BM proteins, into the open circulatory system^11,21,23,34^ and this could play a direct role in the attraction or engulfment of the larval epidermal cells. Persistence of larval epidermal cells may also explain the presence of the semi-transparent, plastic-like sheath surrounding the abdomen of *Tub*>*Glt-RNAi* pharate adults. This sheath and incomplete pupal-adult apolysis are the likely causes pharate adults failed to eclose because they remained attached to the pupal case. Larval epithelial cells are largely responsible for secreting the posterior half of the pupal cuticle^35,36^ and we speculate the semi-transparent sheath is superfluous pupal cuticle that accumulated due to the incomplete removal of the larval cells.

Melanization is a common defense mechanism in arthropods that occurs in response to pathogenic infections and tissue damage^37,38^. Melanin formation at the wound site is known to occur in *Drosophila* as part of the systemic wound response^39^. The frequent observation that *Tub*>*Glt-RNAi* flies exhibit melanotic spots and darkened pigmentation may be an indication of underlying tissue damage due to a weakened BM unable to withstand normal physiologic forces such as those from contracting muscle and pulsating hemolymph. This conclusion is further supported by our observation that premature mortality was most common among *Tub*>*Glt-RNAi* adults exhibiting darkened pigmentation. In addition, expression of Glutactin in hemocytes has been shown to contribute to protection against nematobacterial infection of *Drosophila*. Larva in which Glutactin expression was knocked down in hemocytes showed a significantly higher mortality than control larva when infected with *Heterorhabditis bacteriophora*^40^. The inability to secrete Glutactin may hinder the ability of hemocytes to effectively seal the wounds caused by the nematode^40^. We propose that the high mortality among *Tub*>*Glt-RNAi* flies is a combination of a weakened BM unable to withstand internal stresses and an inability to repair damages that may result from these stresses.

Our results showed that Glutactin co-localizes with the heparan sulfate proteoglycan DPerlecan in the BM surrounding larval muscles. While DPerlecan is required for the presence of Glutactin, the reverse is not true as evidenced by the DPerlecan expression in the BM of *Tub*>*Glt-RNAi* larva. This contrast with Collagen IV whose presence is not dependent on DPerlecan^22^. However, presence and correct deposition of Collagen IV depends on Laminin and Integrin, respectively^2,22,41^. Thus, incorporation of Glutactin into the BM appears to require the prior assembly of most major components. Our studies also raise the possibility that Glutactin is a DPerlecan ligand. Perlecan is known to interact with multiple proteins, including Dystroglycan, Laminin, and growth factors, among others. A possible interaction with Perlecan also explains the multiplicity of phenotypes manifested by *Tub*>*Glt-RNAi* animals given Perlecan’s role in signaling pathways and mechanical events controlling a variety of morphogenetic processes^22,42,43^.

There are no known *in vivo* interactive partners for Glutactin. A previous *in vitro* study that tested ~5,000 individual *Drosophila* proteins using coaffinity purification coupled to mass spectrometry identified 42 proteins interacting with Glutactin^44^. Among these are several extracellular matrix proteins including Collagen type IV (*Cg25c*), Laminin (*LamB2*), and Viking (*vkg*). DPerlecan (*trol*) was not among the proteins included in the study.

The finding that Glutactin is present in larval muscle was unexpected. Its co-localization with phalloidin throughout the thickness of the fiber suggest it is a component of the I band/Z band and not of the costamere, a structure that encircles the Z band and connects the myofibril to the sarcolemma. Furthermore, the slower crawling speed of larva in which Glutactin expression was down-regulated by a mesoderm-specific driver indicates it has a functional role in muscle. However, down regulation of Glutactin had no visible impact on sarcomere registry, though its absence in the DPerlecan hypomorph *trol*^G0271^ coincided with abnormal striations. This is in contrast to the Z-line protein *Thin* whose absence results in sarcomere disorganization and mislocalization of costameric proteins^45^. While we did not examine the effect of Glutactin down-regulation of costameric protein localization, previous studies have shown that Glutactin does not induce costamere formation in *Drosophila* cell cultures^11^. Further studies will be needed to determine if Glutactin contributes to the development and structural organization of sarcomeres and to ascertain its specific role in muscle function.

## 4. Experimental procedures

### 4.1. Fly strains

All *Drosophila* strains were raised at 25°C on standard cornmeal diet. Targeted gene expression experiments using the UAS/GAL4 system were performed at 25°C and 29°C. As drivers, we used the constitutively active *Tubulin-Gal4/TM3* (a gift from A Duttaroy, Howard Univ.), but replacing TM3 with TM6B, *Hu*, *Tb*, *e*, and two mesoderm specific lines: *24B-Gal4 (w[*]; P{w[+mW.hs]=GawB]how[24B]*) (stock # 1767, Bloomington *Drosophila* Stock Center) and Mef2-Gal4 (y^1^ w*; P{GAL4-Mef2.R}3) (stock # 27390, Bloomington *Drosophila* Stock Center). The *Tubulin-Gal4/ TM6B, Hu, Tb, e* fly line was generated by mating *Tubulin-Gal4/TM3* males to females *w*^1118^; *TM3, Sb, e/TM6B, Hu, Tb, e*, and crossing the *Tb*, non-*w*, non-*Sb* progeny.

As responder fly lines we used the *UAS-RNAi-Glt* (stock #15429, Vienna *Drosophila* RNAi Center; herein referred as *Glt-RNAi*) and the *UAS-Glt* (provided by Akinao Nose^12^). In addition we used *w*^1118^, *trol-GFP* (stock # G00022, FlyTrap GFP Protein Trap Database^46^) and the hypomorph *trol*^g0271^ (*P{w[+mC]=lacW}trol[G0271] w[67c23]/FM7c*) (stock # 11848, Bloomington *Drosophila* Stock Center).

To generate a line carrying two transgenes *Glt-RNAi;UAS-Glt*, we first crossed homozygous females *Glt-RNAi* to homozygous males *UAS-Glt* and the male progeny from this cross *w*^1118^; *Glt-RNAi*/+; *UAS-Glt*/+, were crossed to females *w*^1118^; *ap*^Xa^/*Cy*;TM6B, *Tb, e*. Male and female progeny *w*^1118^; *Glt-RNAi*/*Cy*; *UAS-Glt*/TM6B, *Tb, e*, were crossed and *w*^1118^; *Glt-RNAi*/*Glt-RNAi*; *UAS-Glt*/TM6B, *Tb, e*, male and female progeny were crossed to establish the double homozygous stock.

To generate the strain *trol-GFP; Tub/TM6B, Hu, Tb, e* we crossed virgin females *w*^1118^, *trol-GFP* to males TM3, *Sb, e*/*TM6B, Hu, Tb, e*, and the male progeny from this cross *w*^1118^, *trol-GFP*;+/+;+/TM3, *Sb, e* mated to *w*^1118^, *trol-GFP*;+/+;+/*TM6B, Hu, Tb, e*, females. Male progeny *w*^1118^, *trol-GFP*;+/+;TM3, Sb, *e/TM6B, Hu, Tb, e*, were crossed to females *trol-GFP*;+/+;+/*Tub-Gal4* obtained from a separate cross of females *w*^1118^, *trol-GFP* with males Tub-Gal4/*TM6B, Hu, Tb, e*. From the progeny of this last cross we selected for males and females having the *TM6B, Hu, Tb, e*, balancer and the darkest eye intensity to establish the stock *w*^1118^, *trol-GFP/w*^1118^, *trol-GFP;* +/+; *Tub-Gal4/TM6B, Hu, Tb, e*.

To generate the *w*^1118^, *trol-GFP; Glt-RNAi* line we performed several crosses, the first using males *w*^1118^, *trol-GFP* (stock # G00022) crossed to homozygous females *Glt-RNAi* and selected for virgin females *w*^1118^, *trol-GFP*/+;*Glt-RNAi* /+. For a second cross we used virgins females *w*^1118^, *trol-GFP* crossed to homozygous males *Glt-RNAi* and selected for males *w*^1118^, *trol-GFP; Glt-RNAi*/+. These males were crossed to virgin females *w*^1118^, *trol-GFP*/+; *Glt-RNAi*/+ and progeny with the most intense eye color were backcrossed for several generations.

The stock *trol-GFP; Tub*>*Glt-RNAi* was generated by crossing *w*^1118^, *trol-GFP*; +/+; *Tub-Gal4*/ *TM6B, Hu, Tb, e*, and *w*^1118^, *trol-GFP;Glt-RNAi*.

### 4.2. Larva dissections and immunostaining

Third instar wandering larva were dissected as described by^47^. Briefly, a larva was pinned to Sylgard, dissected in Ca^+2^ free saline (130 mM NaCl, 5 mM KCl, 36 mM sucrose, 5 mM Hepes (pH 7.3), 4 mM MgCl_2_, 0.5 mM EGTA) and fixed in 4% paraformaldehyde in PBS for 30 minutes. Dissected larva were incubated with PBS + 0.5% Triton X-100 for 1 hr, followed by incubation in blocking solution (PBS + 0.5% Triton X-100 + 2% BSA) for 15 minutes and immunostained overnight at 7°C with the following antibodies: mouse monoclonal anti-Glutactin antibody C7906-50 (US Biological, see Supplement 3 for validation of the antibody specificity) diluted 1/250; chicken anti-GFP 1/1000 (Invitrogen A10262); rabbit anti-Obscurin 1/500 (kindly provided by B. Bullard^25^) and rabbit anti-DPerlecan V 1/1500 (kindly provided by S. Baumgartner^20^) in blocking solution. Next day, four washes of 15 minutes each with blocking solution were performed and then secondary antibodies diluted in blocking solution were used overnight at 7°C. Next day, secondary antibodies were removed by four washes using PBS + 0.5% Triton X-100 (15 minutes each). Secondary antibodies used included anti-mouse Alexa Flour 555; anti-chicken Alexa Flour 488; anti-rabbit Alexa Flour 680, 1/1000 (Invitrogen). Actin filaments were stained using 1/20 dilution of a 200U stock solution containing Alexa Flour 647-phalloidin (Invitrogen) for two hours and washed with PBS + 0.5% Triton X-100. Dissected larva were mounted on glass slide using Vectashield H-100 (Vector Laboratories) & coverslip and larva body wall musculature was examined using a Zeiss LSM 510 META Confocal Laser Microscope.

### 4.3. Adults frozen sections and immunostaining

We followed the procedure described previously^48^. Briefly, three to five days old flies were embedded in OCT compound (Tissue-Tek 4583) for sectioning in a microtome cryostat. Adult sections (10 μm) were fixed for 30 minutes in 2% formaldehyde solution and washed three times (5 minutes each) with TBS (10 mM Tris, pH 7.5; 130 mM NaCl; 5 mM KCl; 5 mM NaN3; and 1mM EGTA). The sections were pre-incubated for 30 minutes in blocking solution of 3% BSA in Balanced Salt Solution (BSS: 0.03 M NaCl; 0.05 M KCl; 0.12M MgSO_4_-7H_2_O; 0.005 M CaCl_2_-2H_2_O; 0.01 M Tricine; 0.02 M Glucose; 0.006 M Sucrose; 0.005 M NaN_3_) and then incubated for 1 hr with anti-Glutactin primary mouse monoclonal antibody (US Biological, C7906-50) diluted 1/200. For control sections, mouse monoclonal antibody was substituted with mouse pre-immune serum. After washes, sections were incubated in secondary rabbit anti-mouse Alexa Flour 568 (Molecular Probes) diluted 1/200. Nuclear DNA was stained using DAPI diluted 1/400. Sections were mounted with Vectashield H-100 (Vector Laboratories) and coverslip, then examined using a Zeiss LSM 510 META Confocal Laser Microscope.

### 4.4. Larva crawling assay

Crawling speed of individual third instar wandering larva were determined at 25°C as described previously^17^, with the following modifications. The larva were removed from the wall of the vial using a moistened paintbrush and cleaned gently using a damped kimwipe to remove attached food. Cleaned larva were placed in the center of an evenly illuminated 100 mm x 15 mm molasses plates (25 g food agar, 100 ml molasses, 7.5 ml propionic acid, per liter) and allowed to adapt to the molasses environment and illumination for 30 seconds before performing the assay. At the start of the crawling assay, the larva were placed in the center of the plate and allowed to crawl for one minute or until it reached the edge of the plate. We performed one assay per larva for a total of 30 larva and each larva path was traced on a piece of paper and digitalized in jpeg format. To calculate the crawling speed, digitalized images were analyzed using image J (http://rsb.info.nih.gov/ij/). Statistical analyses were done as described below.

### 4.5. Lifespan studies

Experiments were performed at two different temperatures, 25°C and 29°C. Newly eclosed (< 1 day old) adults (*Tub*>*Glt-RNAi*, *Tub*>/+, and *Glt-RNAi*/+) were collected under CO_2_ for a period of two weeks and placed in standard cornmeal medium vials at a density of 20 flies per vial. Fly vials were incubated on a 12:12 hr on/off light cycle. Vials were examined every 22-to-24 hrs, the number of dead flies recorded and live flies transferred to new vials every 48 hrs.

### 4.6. Egg-Laying Assay

Two to three day-old *Glt-RNAi*, *Tub-Gal4*, and *24B-Gal4* virgin females were placed in individual standard cornmeal medium vials maintained in a 12:12 hr on/off light cycle at 29°C to determine the number of eggs laid. Additionally, *Glt-RNAi* virgin females were mated separately to *Tub-Gal4* and *24B-Gal4* males at 25°C. Virgin female progeny from these crosses (*Tub*>*Glt-RNAi*, and *24B*>*Glt-RNAi*) were placed in individual vials as above. Flies were transferred to new vials every 22-to-24 hrs for a period of two weeks, and the eggs in the old vial were counted. Statistical analyses were done as described below.

### 4.7. Eclosion and mortality studies

Food bottles seeded with the same number of males and females, ~30 each per bottle, were allowed to lay eggs for 4-5 days after which adults were removed and progeny allowed to develop. To record pupal and eclosion mortality, we used bottles from which no adults had eclosed in at least five days. Plastic food bottles were cut in half, avoiding damage to any pupa. Pupal cases that were not empty were classified as early pupa or late pupa based on the absence or presence of eye pigmentation, respectively. Mortality at the pharate adult stage included flies unable to fully eclose and those that fully eclosed but remained attached to the pupal case.

### 4.8. Statistical Analysis and Graphs

All values are means + SE. Differences between groups were examined by one-way ANOVA test followed by a post hoc LSD test (IBM SPSS Statistics Package, v.20.0, Chicago, IL), with values considered significant at p<0.05. Line graph were generated using Plotly (https://plot.ly/) and bar graph generated using GraphPad Prism (v.6.03, GraphPad Software Inc, La Jolla, CA).

## Acknowledgements

This work was supported by NSF grants MCB 1050834, IOB 0718417, and Breazzano Family Endowment funds to J.O.V. P.A.O. was partly supported by NSF 0450339 AGEP. Mass spectrometry was conducted at the Vermont Genetics Network Proteomics Facility, supported by an Institutional Development Award (IDeA) from the National Institute of General Medical Sciences of the National Institutes of Health under grant number P20GM103449. We thank Stefan Baumgartner and Belinda Bullard for providing antibodies and the Bloomington *Drosophila* Stock Center, Vienna *Drosophila* RNAi Center, FlyTrap Stocks (Cooley lab at Yale University), Akinao Nose, and Atanu Duttaroy for providing fly lines. Special thanks to Kelci Lanthier for her assistance in the larva crawling assay and Nichole Bishop and Nichole Bouffard (Microscopy Imaging Center, University of Vermont) for their assistant in the confocal microscope. We are grateful to members of the Vigoreaux lab for helpful discussions and suggestions. The contents of this publication are solely the responsibility of the authors and do not necessarily represent the official views of NSF, NIGMS or NIH.

## Author Contributions

P.A.O and J.O.V. conceived the study, analyzed the data and wrote the manuscript. P.A.O., S.G. and R.H. performed the experiments. B.A.B. conducted the mass spectrometry analyses. All authors contributed to the review and editing of the manuscript.

