## Supplemental figures and tables for "Down-regulation of *Drosophila* Glutactin, a cholinesterase-like adhesion molecule of the basement membrane, impairs development, compromises adult function and shortens lifespan"

Pedro Alvarez-Ortiz

Shawna Guillemette

Rachel Humphrey

Bryan A. Ballif

Jim O. Vigoreaux*

**Supplement 1**

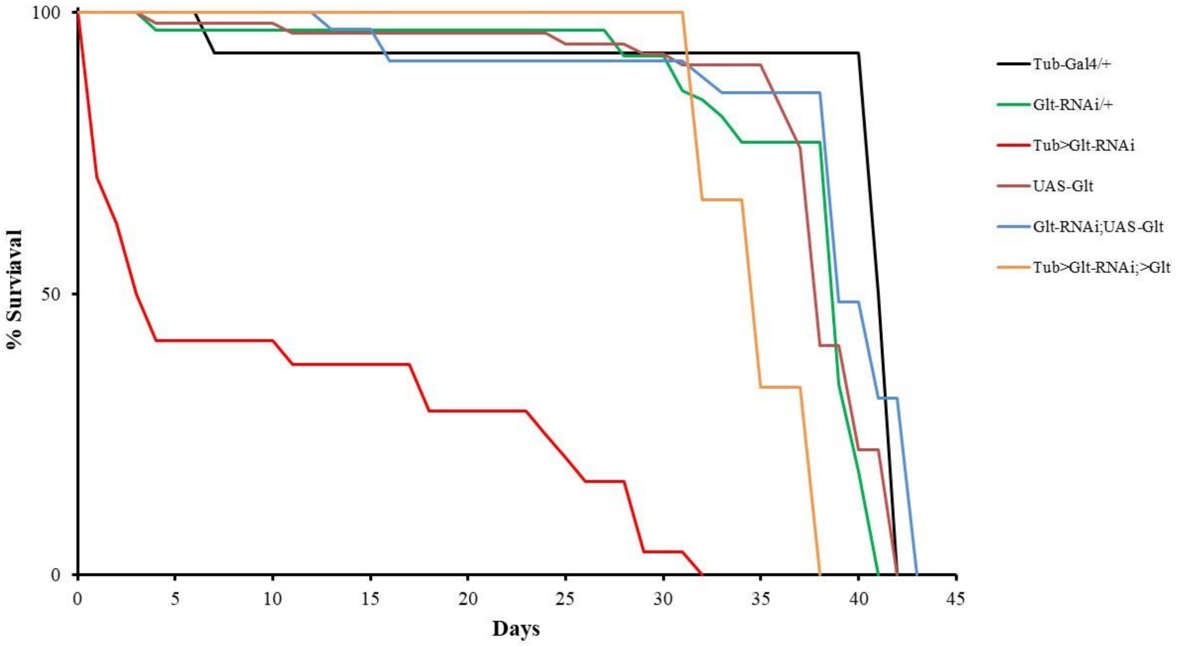


**Figure S1-1. Counteracting effects of over-expression and down-regulation of Glutactin.** Glutactin double mutant (*Tub>Glt-RNAi;>Glt*) circumvent the high mortality brought about by down-regulation of Glutactin (*Tub>Glt-RNAi*). Median lifespan for *Tub>Glt-RNAi;>Glt* (yellow line, n = 3) was 34 days, compared to 3 days for *Tub>Glt-RNAi* (red line, n = 24). Median lifespan for control fly lines were as follows: *Glt-RNAi;UAS-Glt* (blue line, n = 35), 39 days; Tub-Gal4/+ (black line, n = 14), 41 days; *Glt-RNAi/+* (green line, n = 65), 39 days; *UAS-Glt/+* (brown line, n = 54), 38 days. Up-regulation of Glutactin (*Tub>Glt*) resulted in 100% embryonic lethality.

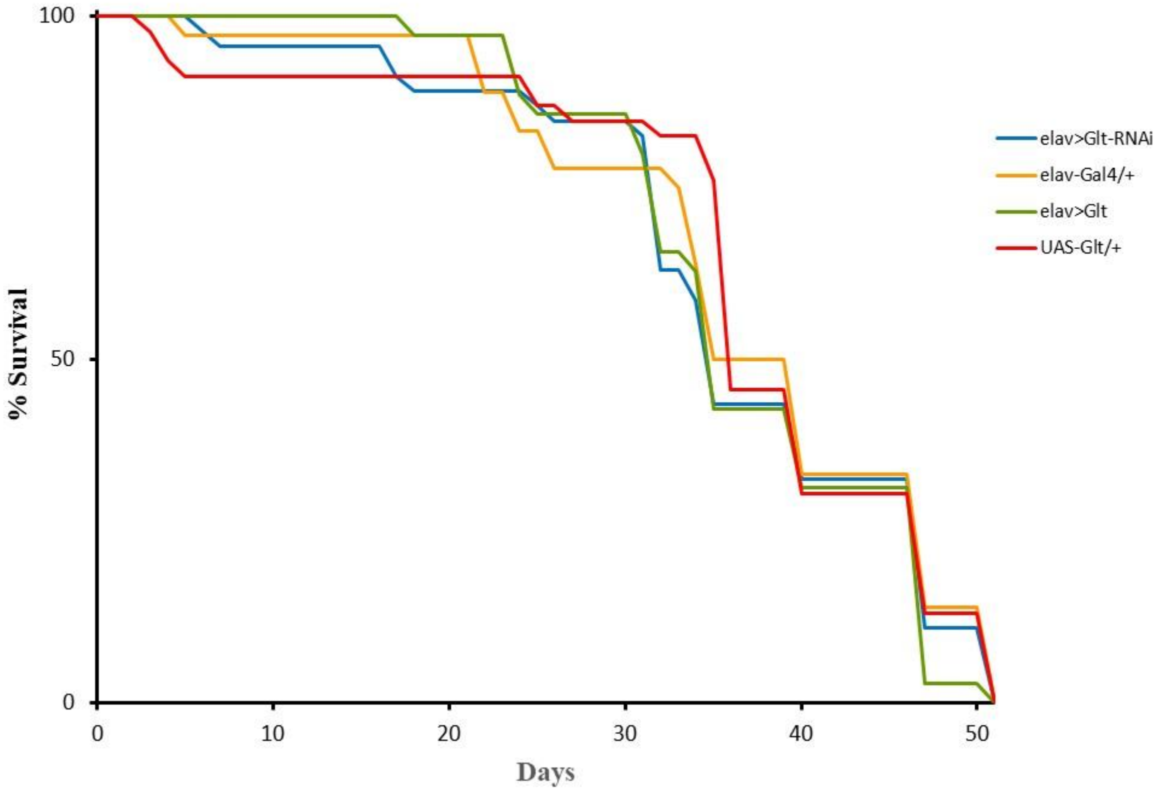


**Figure S1-2. Up-regulation and down-regulation of Glutactin in the nervous system.** Glutactin expression in neuronal cells was controlled using the pan-neuronal Gal4 driver (*elav-Gal4*) (Bloomington Stock 8760). Neither up-regulation (*elav>Glt*, green line, n = 35) nor down-regulation (*elav>Glt-RNAi*, blue line, n = 46) had an effects on lifespan. Also shown are control fly lines: elav-Gal4/+ (orange line, n = 36) and *UAS-Glt/+* (red line, n = 46).

### Supplement 2

**Table S2.1.** Frequency of morphological phenotypes in *Tub>Glt-RNAi* adults.

| Phenotypes | Reference | Frequency (%) 29 $^{o}$C | 25 $^{o}$C |
|---|---|---|---|
| Collapsed body | Fig 2B, 2C | 5 | 0.8 |
| Change in pigmentation | Fig 2E,2F<br>Fig S2B, S2C, S2H, S2I, S2K,S2L | 40 | 26.4 |
| Abdomen dysmorphology | Figure S2E, S2F | 10 | 3.2 |
| Genitalia defects | Figure 2H, 2I, 2K, 2L | 17.5 | 5.6 |
| Impaired locomotion | Supplementary video 1 | 22.5 | 0.85 |

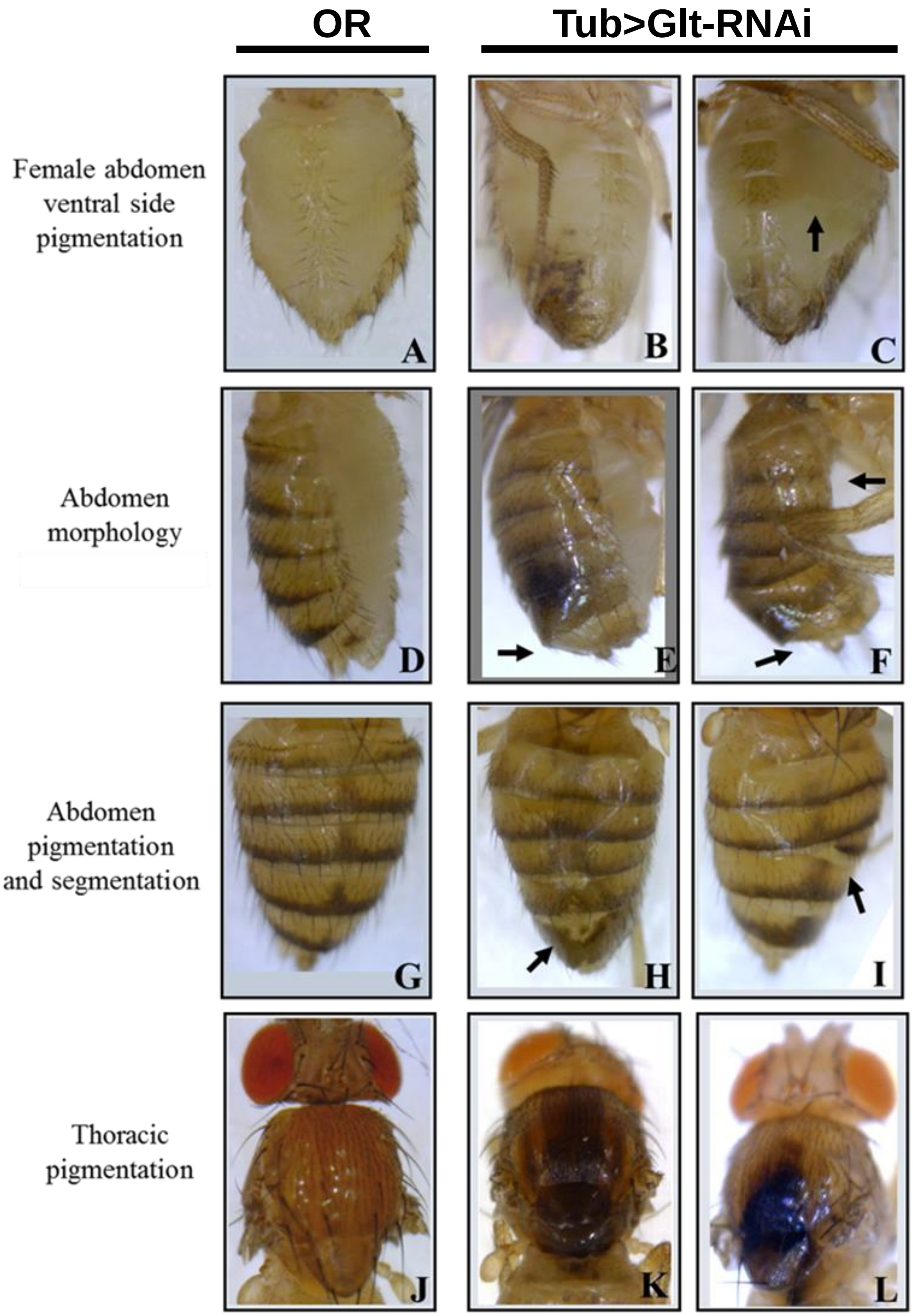


**Figure S2. Morphological changes in adults associated with Glutactin down-regulation by RNAi.** In comparison to other body parts, the abdomen exhibited the most defects as a result of Glutactin down-regulation. Oregon R flies (OR) are shown for direct comparison of the morphological changes (panels A, D, G, and J); all other panels are *Tub>Glt-RNAi* flies. In most panels, wings were clipped to allow better viewing. B and C, examples of female abdomens with necrotic tissue and abnormal pigmentation manifested as a brown shadow (arrow). E and F, examples of flattened posterior end, abdominal segments A7 and A8 appear partially fused. The abdomen in (F) was constricted at the proximal end (right arrow) and enlarged at the distal end (left arrow). The dorsal side of the abdomen exhibits loss of pigmentation (H), and incomplete segment (I). K and L, different shades of thoracic pigmentation.

### Supplement 3

*Validation of mAB C7906-50 specificity*

To validate the use of mouse monoclonal antibody (mAb) C7906-50 (*US Biological) against Glutactin, we performed biochemical and genetic assays to determine its specificity. First, we demonstrate by western blot that mAb C7906-50 recognizes a single *Drosophila* protein band of ~140 kDa present in the larva body wall and in isolated adult thorax, head, and legs, but not in dissected jump muscle and IFM (Figure S3-1). Second, to identify the protein detected by the antibody, we conducted immunoprecipitation of larval proteins with mAb C7906-50 followed by gel electrophoresis and mass spectrometry analysis of immunoprecipitated proteins. Larval proteins were extracted following two different methods; the first was for isolation of extracellular matrix proteins, modified from Fessler et al (Fessler et al. 1994) (Buffer A: 10 mM Triethanolamine, 1% Nonidet P40, 0.5%(w/v) dioxycholate, 0.15 M NaCl, pH 8.2 and 3 μl/ml of protease inhibitor cocktail [CalBiochem Cat# 539134]), and the second for isolation of membrane proteins, using a Zwitterionic detergent (Buffer B: 1.5% Empigen BB [CalBiochem], 1.0% non-detergent sulfobetaines [NDSB] [CalBiochem], 134 mM NaCl, 10 mM phosphate buffer, pH7.4, and 3 μl/ml protease inhibitor cocktail [CalBiochem Cat# 539134]). Briefly, two separate tubes each containing seven 3$^{rd}$ instar larva (OR strain) were frozen in liquid nitrogen, grounded to powder in a mortar and solubilized with either buffer A or B. After adding the buffer, the samples were sonicated with two 15 second pulses. Solubilized mixtures were placed on a nutator for 1 hr at 4$^{o}$C and then centrifuged at full speed for 30 min at 4$^{o}$C. Supernatants were collected and centrifuged twice to remove insoluble material. From each supernatant, 500 μl were used for the immunoprecipitation assay.

Immunoprecipitation assay was performed by adding 30 μl (~90 μg) of mouse anti-chicken cMyBP-C antibodies (US Biological, C7906-50) to each independent extraction tube and to its respective control we added the same volume of solubilization buffer. Each tube was incubated overnight at 4$^{o}$C on a nutator. Next day, 50 μl of beads (Affi-Gel protein A MAPS II kit, Bio-Rad 153-6159) and 580 μl of binding buffer (Affi-Gel protein A MAPS II kit, Bio-Rad) were added to each tube and incubated overnight at 4$^{o}$C on a nutator. Next day, the beads were washed 5 times with 1 ml binding buffer and after the last wash most of the binding buffer was removed. Beads were diluted in 30 µl of 2X LSB (125 mM Tris-HCl (pH 6.8), 0.2 M DTT, 4% SDS, 0.2% bromophenol blue, and 20% glycerol) and then 12 µl were heat up to 95$^{o}$C /5 minutes then loaded in 7% SDS-PAGE followed by staining with SYPRO-RUBY. A prominent protein band of ~140 kDa was detected in the two experimental lanes but absent in each of the corresponding control lanes (Figure S3-2A). The bands were cut out and processed by in-gel digestion and analyzed by mass spectrometry.

In-gel digestion was performed by taking each gel band and dicing it into 1 mm cubes. The cubes were washed with HPLC-grade water, and partially-dehydrated with 50 mM ammonium bicarbonate/50% acetonitrile (MeCN). The cubes were then fully dehydrated with 100% MeCN and allowed to swell on ice for twenty minutes with sequencing grade modified trypsin (Promega, 6 ng/µL). The digestion was then allowed to proceed overnight at 37 °C. Peptides were extracted with 50% MeCN and 2.5% formic acid (FA) and then dried. Peptides were then resuspended in 2.5% MeCN and 2.5% FA and loaded using a Micro AS autosampler (Thermo Electron) onto a microcapillary column packed with 12 cm of reversed-phase MagicC18 material (5 µm, 200 Å, Michrom Bioresources, Inc.). Elution was performed with a 5−35% MeCN (0.1% FA) gradient over 60 min, after a 15 min isocratic loading at 2.5% MeCN and 0.5% FA. The gradient was controlled by a Survey PumpPlus HPLC (Thermo Electron). Mass spectra were acquired in an LTQ-Orbitrap linear ion trap-orbitrap hybrid mass spectrometer (Thermo Electron) over the entire 75 min using 10 MS/MS scans following each survey scan. Raw data were searched against an NCBI non-redundant protein database using SEQUEST software requiring tryptic peptide matches with a 30 PPM mass tolerance. Cysteine residues were required to have a static increase in 71.0371 Da for acrylamide adduction. Differential modification of 15.9949 Da on methionine residues and two internal missed cleavages were permitted.

The top protein hit was Glutactin and was identified by five independent MS/MS scans (Figure S3-2B). Lastly, to validate the mass spectrometry results we used a genetic approach using the *Drosophila* UAS/GAL4 system to down-regulate Glutactin expression by RNA-interference with the constitutively active tubulin-Gal4 line. The ~140 kDa protein band detected in control samples with mAB C7906-50 is absent in the sample from the *Tub>Glt-RNAi* down-regulated line (Figure S3-3).

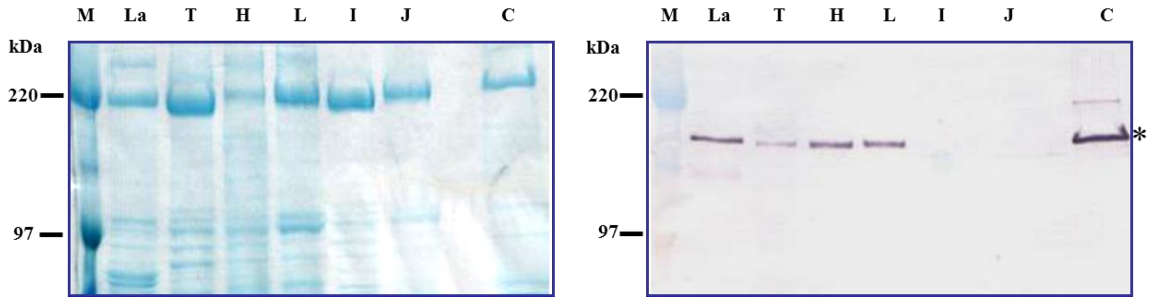


**Figure S3-1. mAb C7906-50 recognizes a *Drosophila* protein band of ~140 kDa.** *Drosophila* proteins from dissected body parts and muscles were separated on a 10% SDS-PAGE and stained with Coomassie blue (left panel) or blotted onto nitrocellulose and probed with mAb C7906-50 (right panel). Lane C corresponds to proteins extracted from chicken cardiac muscle. The band at ~140 kDa (asterisk) is cardiac MyBP-C, shown previously to react with mAb C7906-50 (US Biologicals, Antibody Technical Data). Note that a protein of approximately the same molecular weight as cardiac MyBP-C is detected in *Drosophila* samples from third instar larva body wall (La), whole thorax (T), head (H), and legs (L), but absent in IFM (I) and jump muscle (J). (M) Molecular weight markers, numbers in kilodaltons (kDa). Sample preparation, gel electrophoresis, and western blotting were done as described in (Vigoreaux et al. 1991).

A

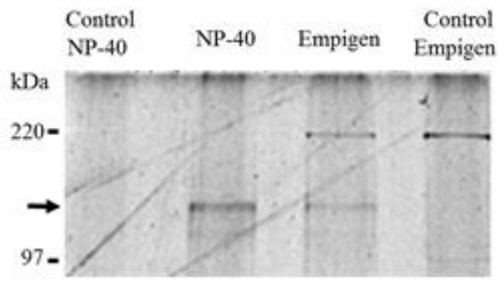


B

| z | XCorr | ΔCorr | Peptide Sequence |
|---|---|---|---|
| 3 | 3.8305 | 0.5862 | R.YAQPQEAVVQAPEVGQILGISGHK.T |
| 3 | 3.5243 | 0.295 | K.LREEQEEQEKLEEER.Y |
| 2 | 3.0261 | 0.3576 | K.DASAFIVSR.I |
| 2 | 2.6272 | 0.2563 | R.IYDQLAR.L |
| 2 | 2.1208 | 0.2102 | R.IYDQLAR.L |

**Figure S3-2. Larval protein immunoprecipitation and peptides identified by mass spectrometry.** (A) 7% SDS-PAGE stained with SYPRO-Ruby (Bio-Rad cat# 170-3125). Whole larva was extracted with two different detergents, NP40 (a non-ionic detergent) and Empigen (a zwitterionic detergent). The two center lanes, labeled NP40 and Empigen, contain the proteins that co-immunoprecipitated with mAb C7906-50. The flanking lanes are the corresponding controls that did not include antibody in the precipitation. The arrow points to a band of ~145 kDa present in the two experimental samples but not detected in the control samples. For each lane the area in the gel that is marked by the arrow was cut out and the proteins were in-gel digested and extracted for mass spectrometry. (B) *Drosophila melanogaster* Glutactin peptides identified by mass spectrometry in the NP-40 and Empigen samples but absent in the control samples are provided with their charge states (*z*) and their SEQUEST XCorr and ΔCn values. The site of tryptic cleavage is denoted by a period.

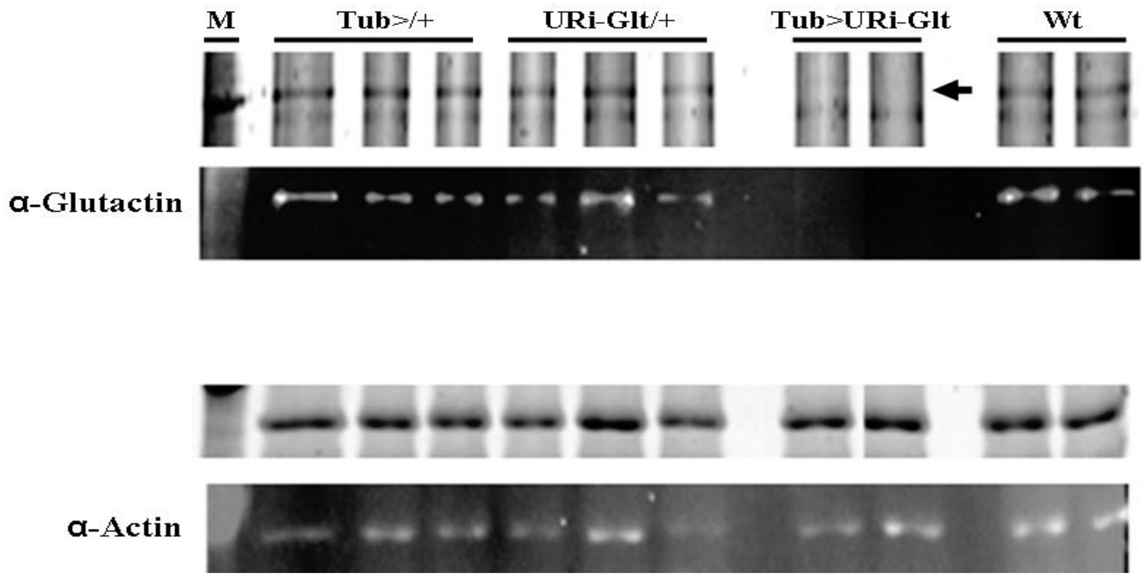


**Figure S3-3. Validation of the detection of Glutactin by mAb C7906-50 in larva.** Glutactin expression is down-regulated by RNAi**.** Body wall proteins from dissected 3rd instar larva of the indicated genotypes were separated on 10% SDS-PAGE and stained with Krypton (Pierce cat# 53072) (top panels) or blotted onto nitrocellulose and probed with monoclonal mAb C7906-50 (Glutactin panel) and polyclonal rabbit anti-actin A-2066 Sigma (actin panel) antibodies. Arrow in top panel points to protein band of ~140 kDa that is not detected in *Tub>Glt-RNAi* lanes. Actin gel and blot are shown as loading controls. See main text for a description of the fly strains.

### Supplement 4

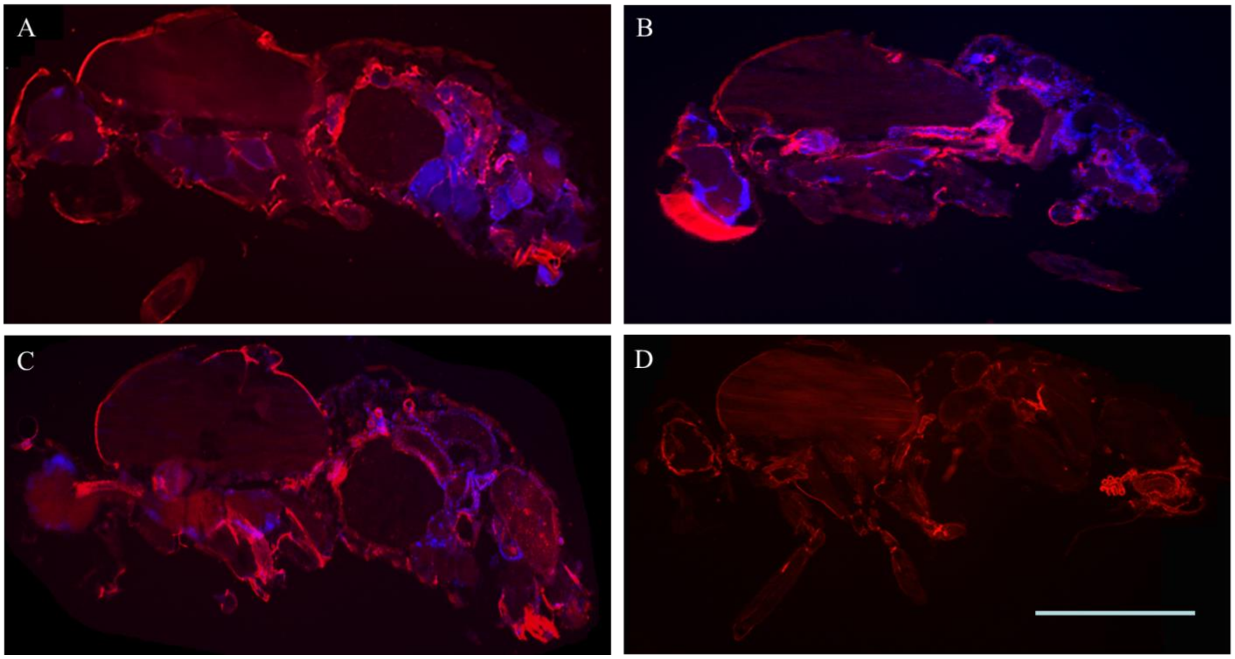


**Figure S4.1. Detection of Glutactin in adults.** All panels are sagittal sections, anterior is to the left and dorsal is up. Panel A-C**:** Adult male double immunostained with anti-Glutactin (red) and DAPI (blue). Panel A and C are sections from the same fly. Glutactin expression is prominent in the abdomen, around the gut, genitalia (A) and rectal ampulla (C). Strong staining is also detected along the head and thoracic cuticle (autofluorescence), the legs, and the alimentary canal (foregut toward the head, which include the esophagus). Panel B, expression of Glutactin in the foregut, proventriculus, and beginning of the midgut. There is strong autofluorescence from the eye. Panel D, distribution of Glutactin in female is similar to that in males. Note the high expression of Glutactin in the reproductive system (posterior end). Scale bar = 1 mm.

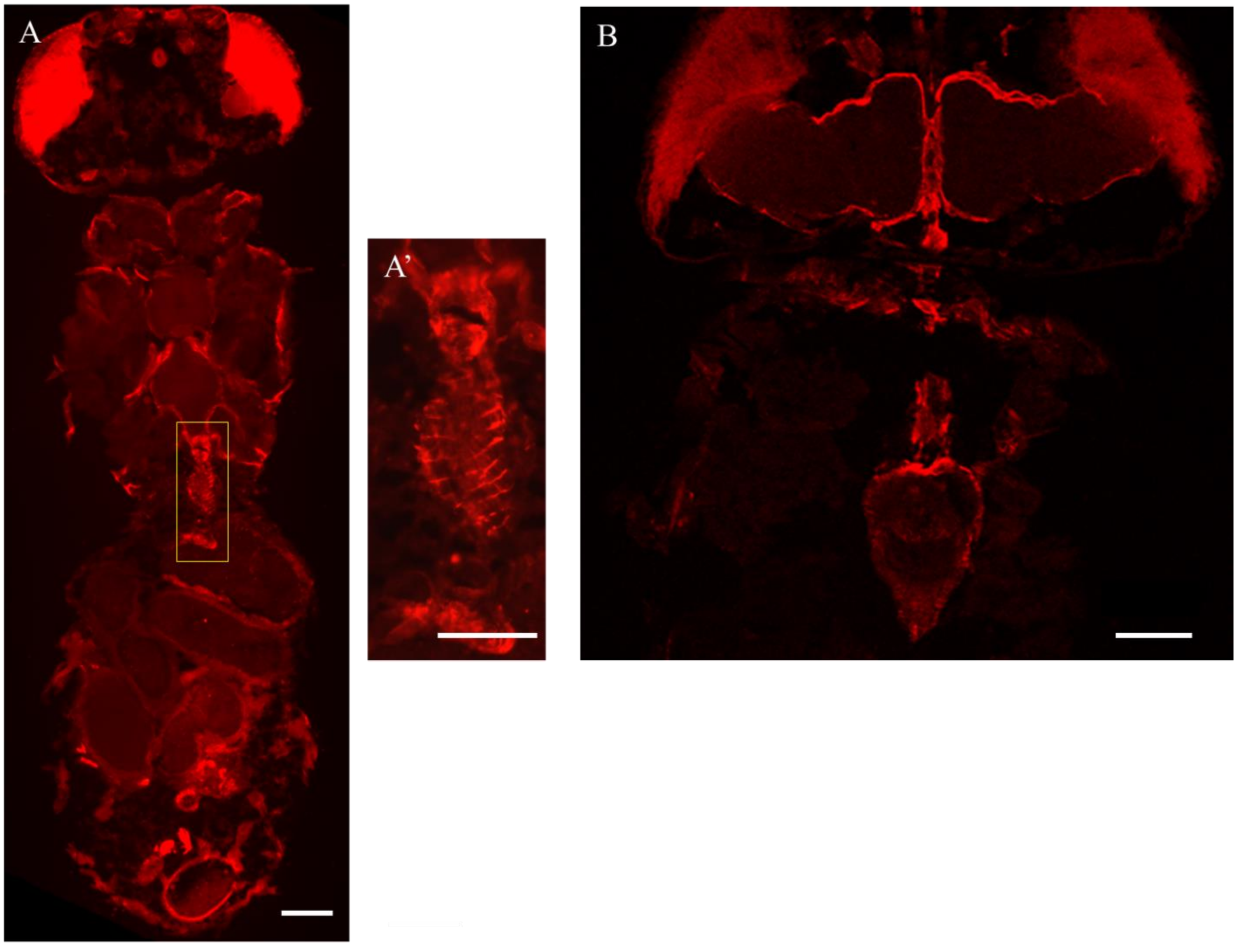


**Figure S4.2. Detection of Glutactin in adult head and thorax.** All panels show dorsal view of female, anterior end is up. Panel A, staining is detected surrounding major thoracic organs, the flight muscle and the foregut. The structure within the yellow square, corresponding to the R3 region of the gut, is shown in higher magnification on the panel A'. Panel B shows a higher magnification of the head and thorax region. Glutactin is detected surrounding the brain and the foregut with strong staining of the heart-shape proventriculus. There is strong autofluorescence from the eyes. Scale bar in Panel A and B = 100 µm, A' = 50 µm.

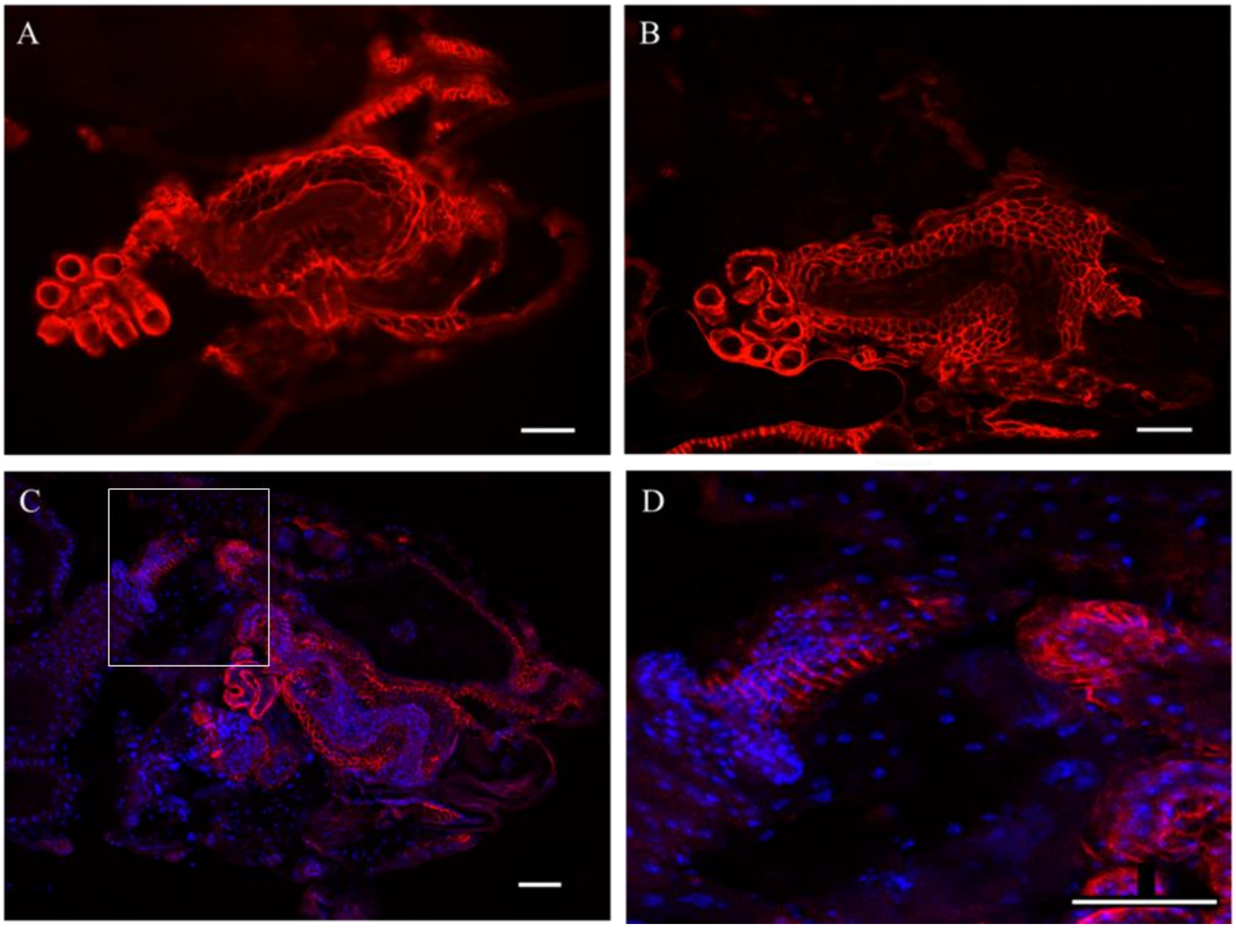


**Figure S4.3. Detection of Glutactin in female reproductive system and abdominal structures.** All panels are adult female sagittal sections, anterior is to the left and dorsal is up. Panel A and B**,** stained with anti-Glutactin (red) and Panel C and D, double immunostained with anti-Glutactin (red) and DAPI (blue). Panel A-C is the same female reproductive organ sectioned at different depth. Panel A-B, left side, note the presence of Glutactin surrounding each ovariole (represented by eight tubes coming out of the picture), which are connected at the base by the extending lateral oviduct (not shown) which connect to the uterus (long wide area) via the common oviduct (not shown). Panel C, lower magnification view of the abdomen showing the same organ as Panel A-B, where we can appreciate Glutactin mostly enriched in the outer layer of cells which are surrounding this organ. In this panel we can also appreciate the detection of Glutactin surrounding the rectal ampulla and rectum (above the reproductive organ having a wide open unstained space). Panel D: higher magnification of boxed region in panel C. Glutactin is detected surrounding the hindgut in a pattern resembling striated muscle. Scale bars in panels A-D = 50 μm.

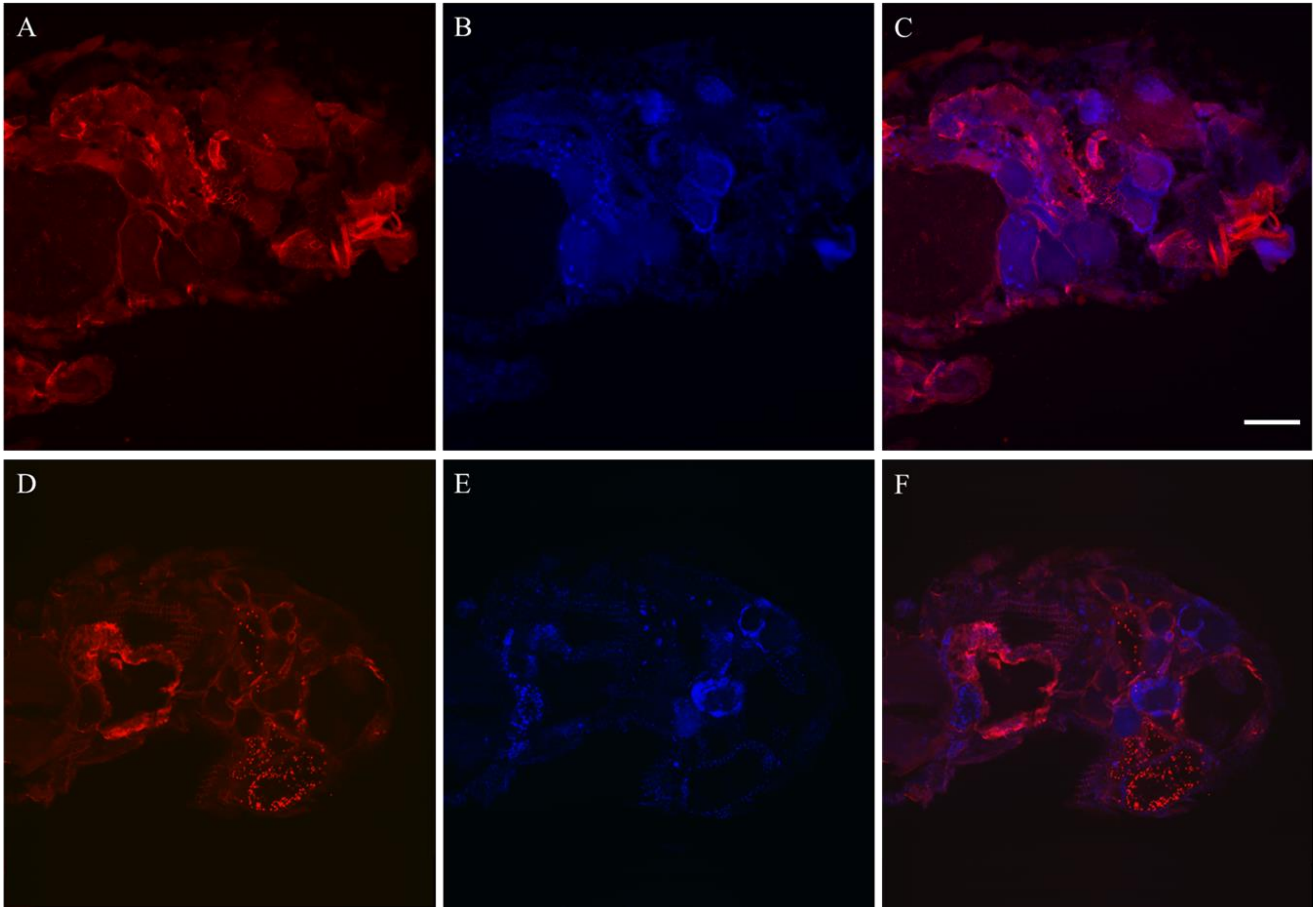
A
B
C
D
E
F

**Figure S4.4. Detection of Glutactin in male reproductive system and abdominal structures.** Sagittal sections of male abdomen, anterior is to the left and dorsal is up. Double immunostaining with anti-Glutactin (red) (Panel A and D) and DAPI (blue) (Panel B and E). Panels C and F show the merge images of A and B, and D and E, respectively. Panels A-C and Panel D-F represent two different adults. Panel A and D, strong staining in ventral-posterior region of the abdomen where testis and gonads are located. Strongest staining in A and C co-localizes with the ejaculatory bulb and the penis. Glutactin is also detected surrounding the gut. Scale bar = 100 μm.

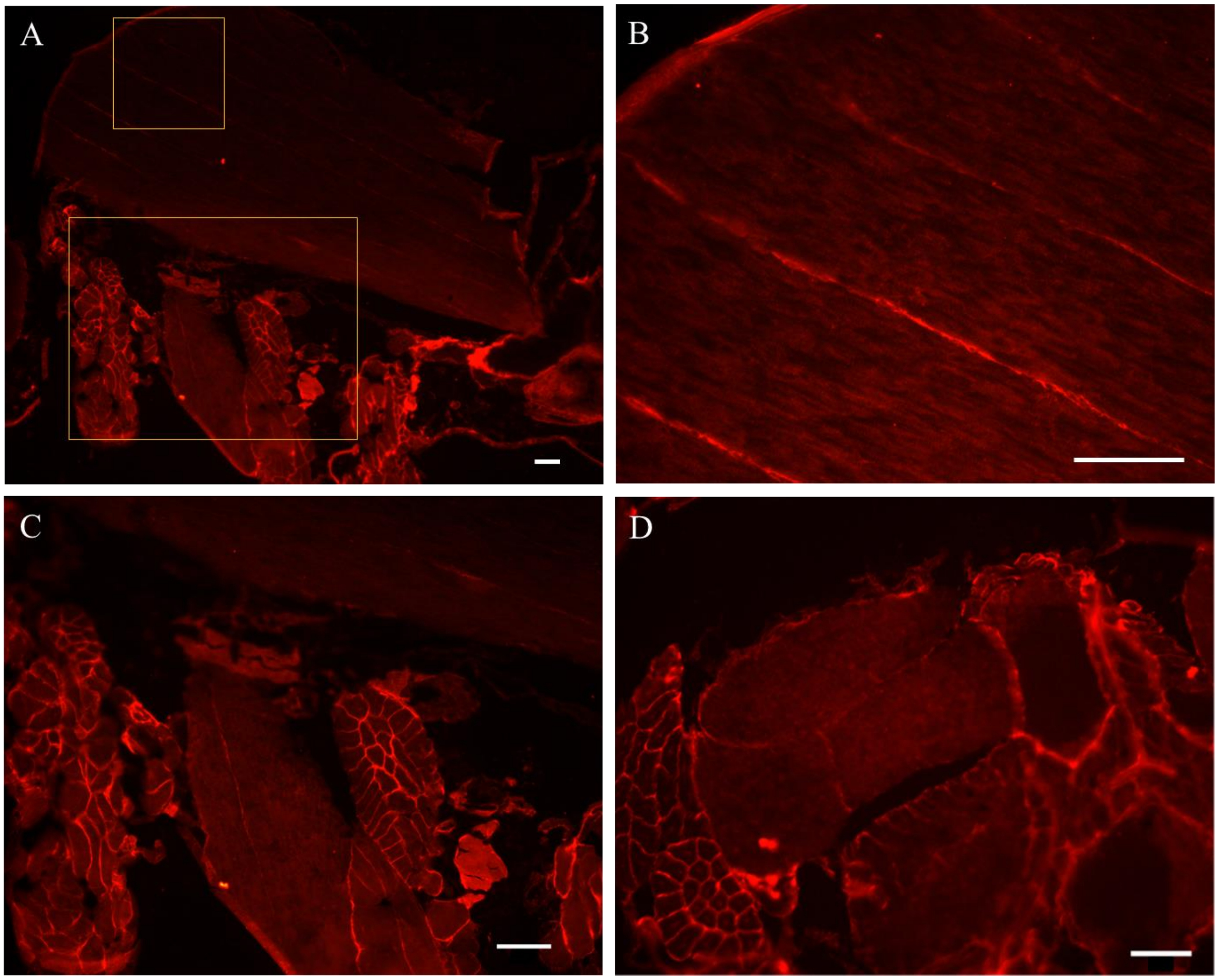


**Figure S4.5. Detection of Glutactin in the thorax.** Panel A-C**,** sagittal section of the thorax showing the dorsal longitudinal indirect flight muscles (fibers extending from top left to bottom right) and legs muscles just below them. Ventral side is down. The two areas demarcated by the yellow square are shown in higher magnification in Panels B and C. Panel B, note the presence of Glutactin in the interstices between the muscle fibers. Panel C, note Glutactin surrounding the tubular muscles of the legs. Panel D, dorsal view of flight muscles in the thorax. Glutactin is detected surrounding the direct flight muscles (far left side) and the three big bundles of the dorsal longitudinal indirect flight muscles. For all images, anterior is to the left. Scale bars in panels A-D = 50 μm.

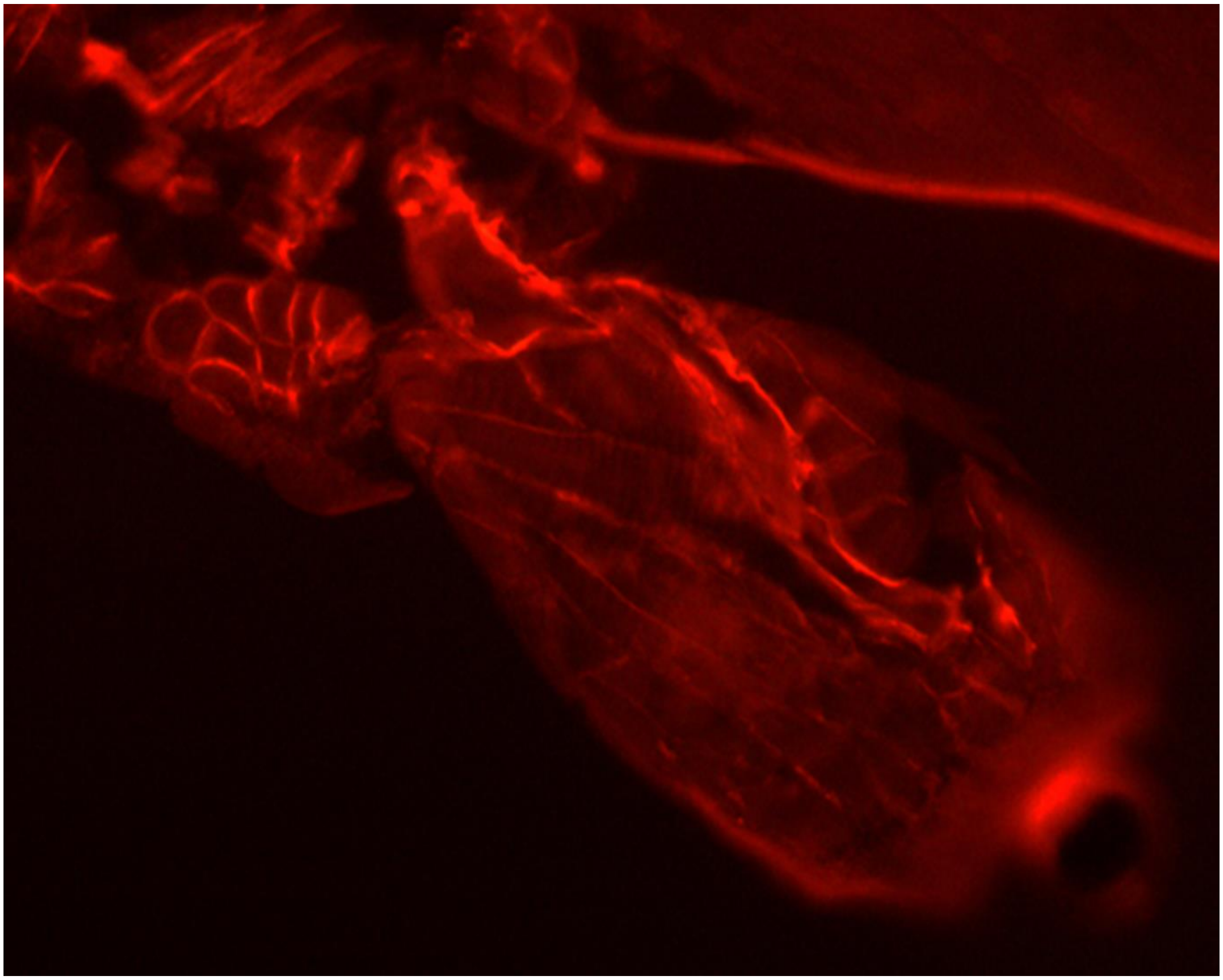

**Figure S4.6. Detection of Glutactin in the legs.** Longitudinal section of the coxa showing the presence of Glutactin surrounding tubular muscles and in a repetitive pattern resembling muscle striations.

### Supplement 5

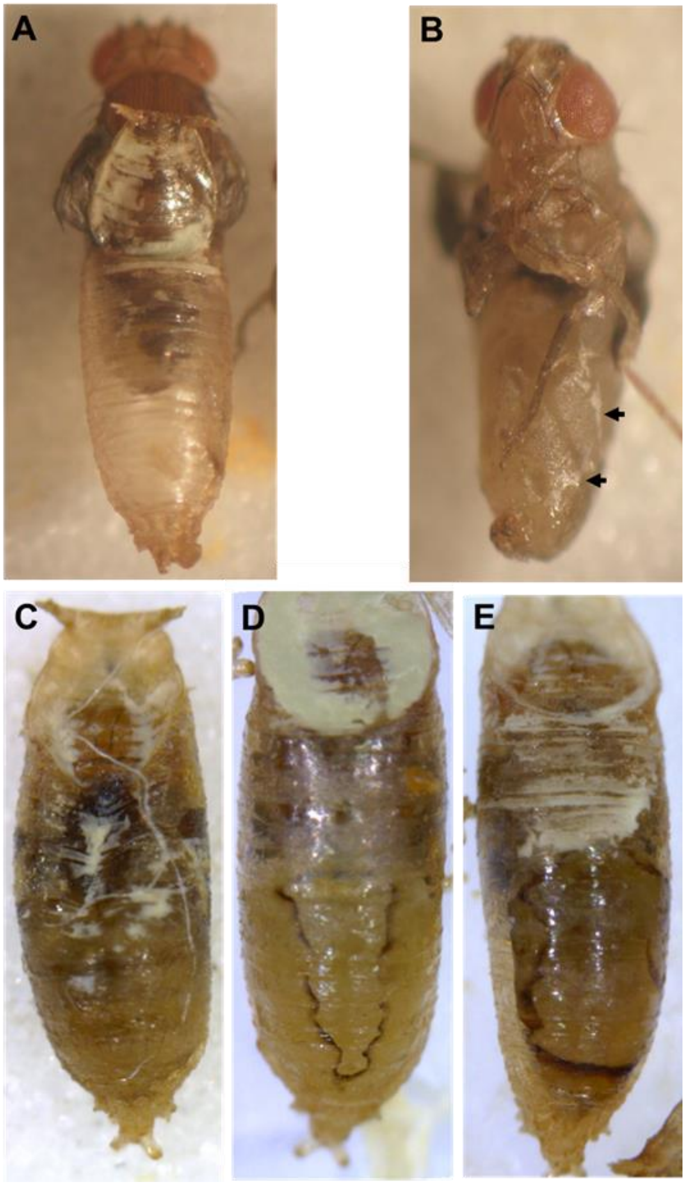


**Figure S5.1. Effects of down-regulation of Glutactin in pharate-adults and pupa.** For all panels, anterior is up. Panel A, C-E are dorsal views while panel B is ventro-lateral. Panel A, a dead pharate-adult with white material on pupal case, similar to what was detected in other dead pupa, panels C-E. Panel B, a dead pharate-adult pulled out from the pupal case to reveal white material (arrows) adhered to the abdomen. Panels C-E, dead pupa showing different amounts of accumulated off-white material in the anterior region. Note in the posterior region of C vs D-E, the region of developing abdomen, varying severity of open scars from none (C) to more (D) and most (E) severe.

### Supplementary Videos

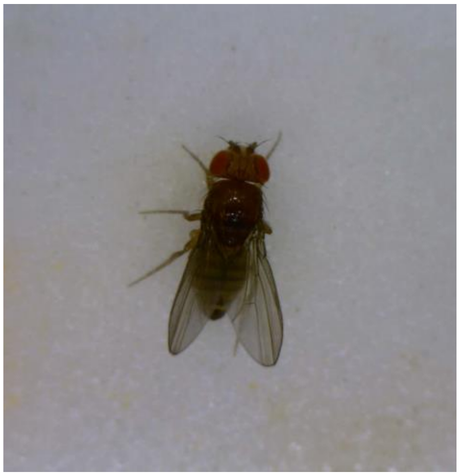

Video 2- 1 .wmv

**Supplementary Video 1.** RNAi down-regulation of Glutactin expression affects adult locomotory functions. One day old adult *Tub>Glt-RNAi*.

Link: https://www.youtube.com/watch?v=qzqZJjas7jQ

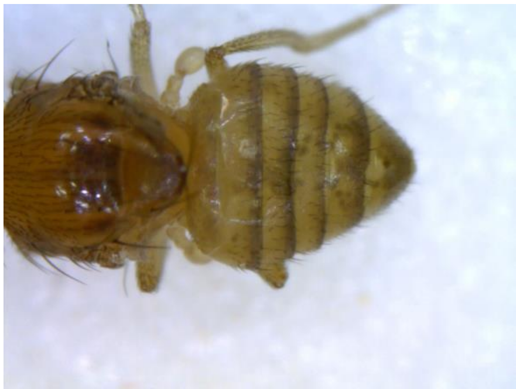

Video 2-2 .wmv

**Supplementary Video 2.** RNAi down-regulation of Glutactin expression affects morphology and contraction of the heart. One day old adult *Tub>Glt-RNAi*.

Link: https://www.youtube.com/watch?v=VQvjUUemZUw
